# Geometry shapes cytoplasmic Cdk1 waves that drive cortical dynamics

**DOI:** 10.64898/2026.03.21.713419

**Authors:** Daniel Cebrián-Lacasa, Marcin Leda, Andrew B. Goryachev, Lendert Gelens

**Affiliations:** Laboratory of Dynamics in Biological Systems, Department of Cellular and Molecular Medicine, KU Leuven, Herestraat, 49, Leuven, Belgium; Centre for Engineering Biology, School of Biological Sciences, University of Edinburgh, C.H. Waddington Building, Max Born Crescent, University of Edinburgh, Edinburgh EH9 3BF, United Kingdom

**Author notes:** **For correspondence:** (LG); (ABG).

## Abstract

Cell division in large embryos is coordinated by spatial waves of Cyclin B–Cdk1 activity that spread through the cytoplasm and drive cortical contractility. However, it remains unclear how cell size and nuclear enrichment of Cyclin B–Cdk1 shape these waves, and how the cytoplasmic signal is transmitted to the cortex. Here we couple established reaction–diffusion models of Cyclin B–Cdk1 signaling and the excitable Rho–actin cortex to ask how cell geometry and elevated nuclear Cyclin B–Cdk1 concentration shape cytoplasmic wave propagation in spherical cells. We find that cytoplasmic waves consist of two distinct parts: an activation front that propagates in a manner consistent with trigger-wave behavior, and a wave back controlled by concentration gradients in the cell cycle oscillator. Because these parts are generated by different mechanisms, they can move at different speeds or even in opposite directions, depending on nuclear size and effective cell size. This yields a simple geometric explanation for why surface contraction waves propagate in opposite directions in starfish oocytes and Xenopus embryos, without invoking system-specific cortical mechanisms. Coupling the Cdk1 signal to the cortex further shows how cytoplasmic waveforms regulate Rho–actin dynamics through inhibition of the Rho GEF Ect2. Together, these results provide a mechanistic framework linking localized nuclear activation, cytoplasmic Cdk1 waves, and cortical responses in large embryonic cells.

## Introduction

Cyclin B–Cdk1 is the core regulator of meiotic and mitotic entry. In large cells, this raises the question of how Cdk1 activity is coordinated across the cytoplasm. Early theoretical work established that positive feedback loops endowing the Cdk1 network with bistability can support propagating trigger waves—self-regenerating activation fronts that advance at constant speed through local positive feedback, rather than by the passive, ever-slowing spread of diffusion—providing a mechanism for such coordination ***Novak and Tyson (1993)***. Indeed, in early embryos and oocytes, where cells can reach hundreds of microns to millimeters in size, Cdk1 activation does not occur uniformly throughout the cytoplasm. Instead, Cdk1 activity propagates as a large-scale cytoplasmic wave following nuclear envelope breakdown (NEB) ***Deneke et al. (2016)***; ***Bischof et al. (2017)***; ***Chang and Ferrell Jr (2013)***; ***Puls et al. (2024)*** (Box 1, Fig. 1). Such waves coordinate cell cycle progression across distances that far exceed typical molecular diffusion lengths and therefore require mechanisms beyond simple diffusive equilibration. More broadly, the spatial coordination of biochemical oscillations via inter- and intracellular waves has emerged as a key focus of research, as wave-driven coordination is critical for maintaining proper cellular function and developmental processes ***Gelens et al. (2014)***; ***Beta and Kruse (2017)***; ***Deneke and Di Talia (2018)***.

**Figure 1.**
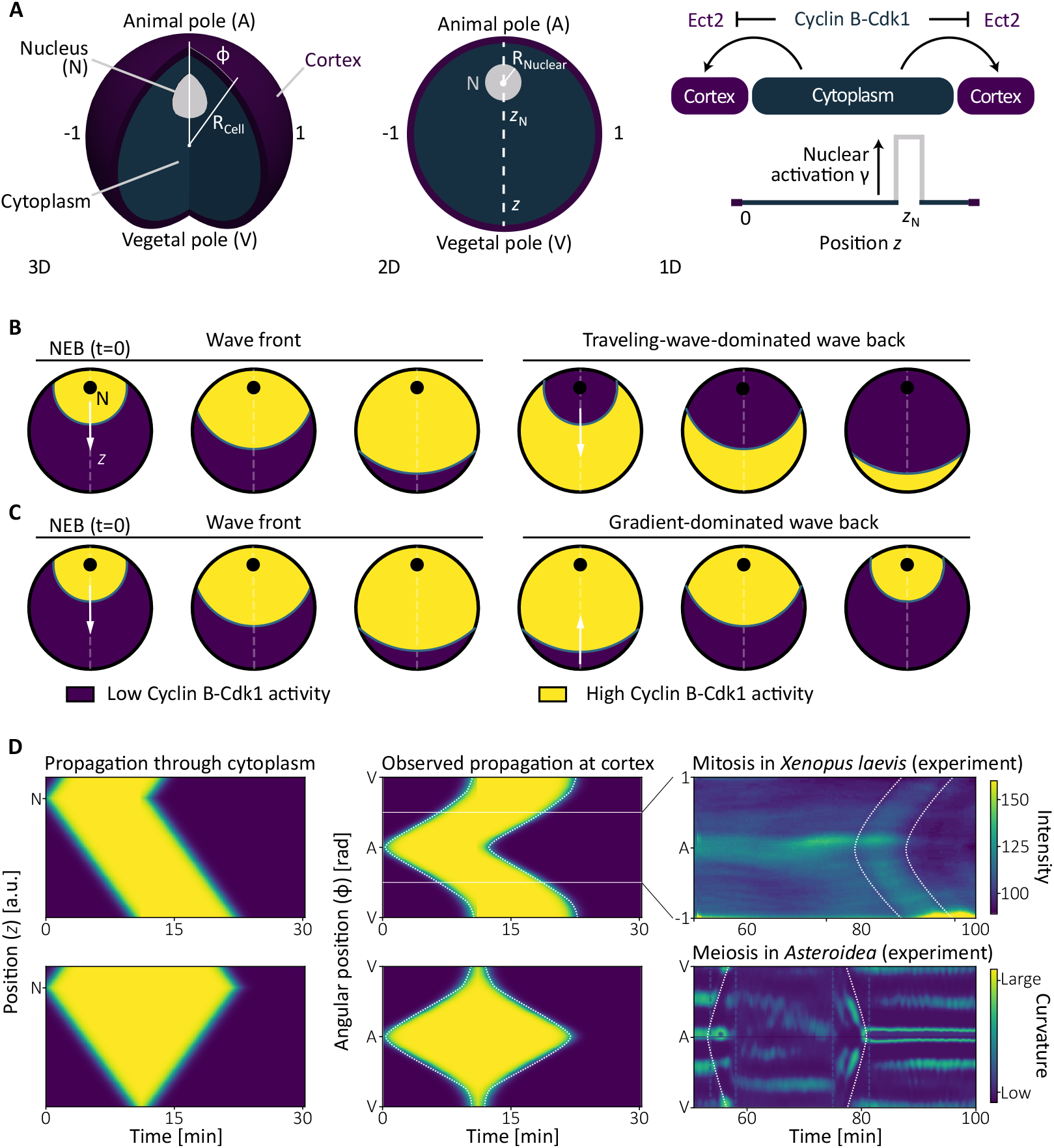
A coupled cytoplasm–cortex model and representations of wave directionality. **A.** Model geometry and coupling. Left (3D) and middle (2D): a spherical cell of radius *R*_Cell_ with an off-center nucleus (radius *R*_Nucleus_) positioned along the animal–vegetal axis at *z_N_*; *z* is the radial coordinate from the nucleus and *φ* the angular coordinate measured from the animal pole. Right (1D): the reduced radial description used throughout. Cytoplasmic Cyclin B–Cdk1 acts as a localized nuclear source, set by the nuclear enrichment factor *γ*, and inhibits the cortical Rho GEF Ect2 at both poles. **B.** Schematic of active Cyclin B–Cdk1 propagation in a traveling-wave–dominated regime, shown as snapshots on the cell (nucleus in black, *z*-axis dashed): an activation front sweeps from the nucleus to the cortex, followed by a wave back that propagates in the same direction. **C.** Same representation for a gradient-dominated regime, in which the wave back reverses and collapses back toward the nucleus. Purple, low Cyclin B–Cdk1 activity; yellow, high activity (shared legend below B,C). **D.** Kymographs of the dynamics sketched in B,C. Left: activity along the cell diameter through the nucleus (the *z*-axis). Middle: the same dynamics along the angular coordinate *φ* defined in A, as seen at the cortex. Right: experimental kymographs from biological systems showing these dynamics; color encodes the plotted quantity (top, Cdk1-reporter intensity; bottom, local membrane curvature), with dotted lines marking the tracked front and back. Top: mitosis in *Xenopus laevis* (replotted from data digitized from ***Chang and Ferrell Jr (2013)***, with additional annotations). Bottom: meiosis in starfish (replotted from data digitized from ***Bischof et al. (2017)***, with additional annotations).

Cytoplasmic Cdk1 waves are directly coupled to the cell cortex ***Foe and Alberts (1983)***. The cortex is a dynamic actomyosin network that controls membrane tension, surface deformation, and cell shape ***Nalbant et al. (2004)***; ***Bement et al. (2005)***; ***Machacek et al. (2009)***. Rho GTPases are central regulators of cortical patterning and their spatiotemporal organization underlies a wide range of cortical behaviors ***Beta et al. (2023)***; ***Bement et al. (2024)***. During meiosis and mitosis, spatial variation in Cdk1 activity is transmitted to the cell cortex through phosphorylation of the Rho GEF Ect2, which reduces its membrane affinity and inhibits Rho GTPase signaling ***Hara et al. (2006)***; ***Niiya et al. (2006)***; ***Su et al. (2011)***. The resulting cortical response manifests as surface contraction waves (SCWs), consisting of a relaxation phase (SCWa) followed by a contraction phase (SCWb), first described by Hara in amphibian eggs ***Hara (1971)*** and later characterized in multiple systems ***Foe and Alberts (1983)***; ***Rankin and Kirschner (1997)***; ***Bischof et al. (2017)***. The enslavement of cortical and bulk actomyosin contractility to Cdk1 activity is particularly well established in *Drosophila* and zebrafish embryos, where it has been linked to concrete functions such as nuclear positioning and ooplasmic segregation ***Deneke et al. (2016)***; ***Shamipour et al. (2019)***. By comparison, the systems we consider here—the meiotic and early mitotic waves of oocytes and large eggs—remain less well understood at the level of how cytoplasmic wave geometry sets the cortical response.

### Box 1.

Cyclin-Cdk1 dynamics and mitotic waves

**Box 1—figure 1.**
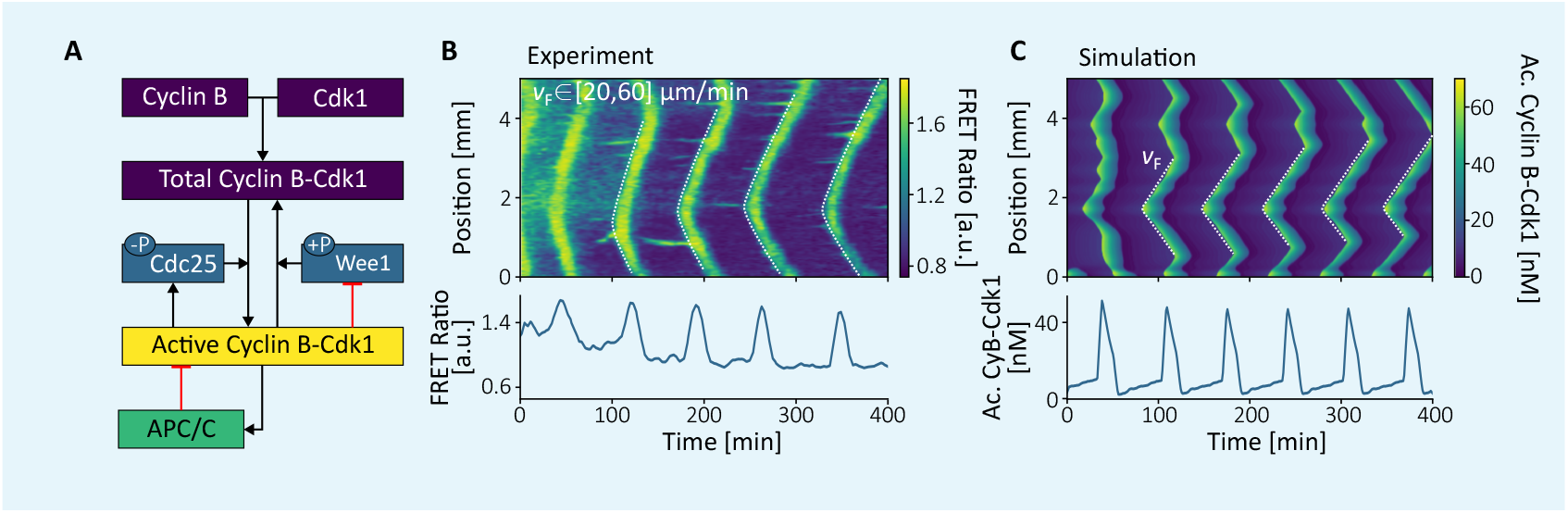
The Cyclin B-Cdk1 network and its dynamics. **A.** Basic network structure regulating the Cyclin B–Cdk1 enzymatic complex. Through phosphorylation and dephosphorylation, Cyclin B–Cdk1 establishes positive feedback loops encompassing Cdc25 and Wee1. In addition, regulation of APC/C generates a delayed negative feedback loop. Together, these features form the essential ingredients for relaxation-like oscillations, characterized by rapid transitions between active and inactive states and robustness to perturbations. **B.** Experimental measurements of Cyclin B–Cdk1 activity in frog cell-free extracts, obtained using a FRET sensor. The time series displays the characteristic oscillations described in panel A. The spatiotemporal dynamics shown in the kymograph further illustrates how, in the presence of nuclei, activity becomes synchronized through traveling waves. Data digitized and replotted from ***Puls et al.*** (2024), with modified axis ranges and color scales. **C.** Computational models have successfully reconstructed the key mechanisms underlying the spatiotemporal dynamics of Cyclin B–Cdk1. In this work, to connect cellular geometry to emergent wave propagation regimes we use the model introduced by ***Yang and Ferrell Jr*** (2013) and extended to a spatial framework by ***Puls et al.*** (2024). Data digitized and replotted from ***Puls et al.*** (2024).

SCWs have been documented in many systems, yet their directionality has been interpreted in different ways. During mitosis, SCWs have been proposed to reduce sensitivity to cell shape during division plane positioning ***Minc and Piel (2012)***, a role for which directionality appears secondary. In contrast, during meiosis, SCW directionality correlates with polar body emission in some organisms. In starfish oocytes, SCW propagation and collapse have been associated with the site of polar body extrusion ***Satoh et al. (2013)***, although subsequent work has shown that cytoplasmic flows generated by cortical contractions are insufficient to reposition the spindle and that polar body emission can still occur when SCWb is inhibited ***Klughammer et al. (2018)***. These findings suggest that SCWs may function primarily as a coordinated cortical response to an underlying cytoplas-mic signal rather than as an autonomous cortical patterning mechanism ***Bement et al. (2024)***; ***Liu et al. (2025)***. A striking illustration of the puzzle this raises is that SCWs run in opposite directions in different systems: they co-propagate towards the vegetal pole in *Xenopus* embryos ***Chang and Ferrell Jr (2013)***, whereas in starfish oocytes a component of the wave collapses back toward the animal pole ***Bischof et al. (2017)***—even though both rely on the same conserved Cyclin B–Cdk1 and Rho–actin machinery. Why the same molecular system should produce opposite cortical behaviors has remained unclear.

Here we show that this difference need not reflect system-specific cortical mechanisms, but can instead follow from cell geometry and nuclear concentration of Cyclin B–Cdk1 alone. Rather than developing a new model, we couple two established reaction–diffusion models—a spatially extended Cyclin B–Cdk1 oscillator ***Yang and Ferrell Jr (2013)***; ***Puls et al. (2024)*** and an excitable Rho–actin cortex ***Michaud et al. (2022)***—in a spherical cell with a localized, off-center nucleus, and couple them through Cdk1-dependent inhibition of the Rho GEF Ect2 (Fig. 1A). Within this geometry we ask how nuclear size, nuclear position, nuclear Cyclin B–Cdk1 content, and effective cell size shape the response. Focusing first on the cytoplasm, we find that Cdk1 waves generically consist of two mechanistically distinct components. Activation fronts propagate with approximately constant velocity, consistent with trigger-wave behavior, whereas wave backs are shaped by diffusion-driven Cdk1 gradients rather than by local bistable kinetics (illustrated schematically in Fig. 1B,C). Because the two components respond differently to nuclear Cyclin B–Cdk1 content and effective system size, they can propagate at different speeds and even in opposite directions, giving rise to a family of wave behaviors within a single conserved network (Fig. 1D). Coupling this cytoplasmic signal to the cortex through Ect2 inhibition, we then show that the resulting SCWs emerge as downstream, phase-wave responses to cytoplasmic signaling geometry and recovery dynamics, providing a simple geometric account of the cross-species directionality difference.

## Model

We couple two established reaction–diffusion models in the spherical geometry sketched in Fig. 1A: a cytoplasmic Cyclin B–Cdk1 oscillator, active throughout the cell but initialized with elevated activity in the nucleus, and an excitable Rho–actin cortex whose distance to a bifurcation is gated by the cytoplasmic signal. The two networks are detailed in Box 1 and Box 2; here we specify each module and the coupling between them.

### Cytoplasmic Cyclin B–Cdk1

The Cyclin B–Cdk1 oscillator is governed by interconnected positive and negative feedback loops that generate sustained oscillations and support trigger wave propagation ***Novak and Tyson (1993)***. We use the heuristic temporal model of ***Yang and Ferrell Jr (2013)***, extended to a spatiotemporal framework with diffusive terms ***Puls et al. (2024)***,

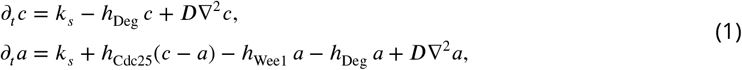

where *a*(**x**, *t*) and *c*(**x**, *t*) denote the active and total Cyclin B–Cdk1 concentrations and *D* is the diffusion coefficient. The ultrasensitive regulatory terms ℎ_Cdc25_, ℎ_Wee1_, and ℎ_Deg_ representing Cdc25, Wee1, and APC/C are specified in the Methods and Materials; all parameters are taken from ***Yang and Ferrell Jr (2013)*** and lie within experimentally measured ranges (see Tab. 1).

**Table 1.** Standard parameter values used in all simulations unless stated otherwise. Cytoplasmic parameters follow ***Yang and Ferrell Jr (2013)***; cortical parameters follow ***Michaud et al. (2022)***. Where four values are given, they are listed as [Regime (i), (ii), (iii), (iv)], corresponding to the four dynamical regimes of Fig. 2C: traveling-wave–dominated, counterpropagating back, coexistence, and homogeneous relaxation. Here *γ* is the nuclear enrichment factor and *s* the inhibition strength. The nuclear radius (*R*_Nucleus_) is used for these four regimes and is varied separately in Supplementary Fig. S3.

| Cortical model |  |  | Cytoplasmic model |  |  |
|---|---|---|---|---|---|
| Parameter = | Value | Units | Parameter = | Value | Units |
| *k*<sub>0</sub> = | 0.00625 | s<sup>−1</sup> | *k*<sub>*s*</sub> = | 1.5/60 | nM s<sup>−1</sup> |
| *k*<sub>1</sub> = | 0.3125 | *µ*M<sup>−3</sup>s<sup>−1</sup> | *a*<sub>Deg</sub> = | 0.01/60 | s<sup>−1</sup> |
| *k*<sub>2</sub> = | 1 | *µ*M<sup>−2</sup> | *b*<sub>Deg</sub> = | 0.06/60 | s<sup>−1</sup> |
| *k*<sub>3</sub> = | 0.0625 | s<sup>−1</sup> | *n*<sub>Deg</sub> = | 17 | – |
| *k*<sub>4</sub> = | 0.05625 | *µ*M<sup>−1</sup>s<sup>−1</sup> | EC<sub>50,Deg</sub> = | 32 | nM |
| *k*<sub>5</sub> = | 0.0625 | *µ*M s<sup>−1</sup> | *a*<sub>Cdc25</sub> = | 0.8/60 | s<sup>−1</sup> |
| *k*<sub>6</sub> = | 0.02083 | s<sup>−1</sup> | *b*<sub>Cdc25</sub> = | 4/60 | s<sup>−1</sup> |
| *k*<sub>7</sub> = | 0.001875 | *µ*M s<sup>−1</sup> | *n*<sub>Cdc25</sub> = | 11 | – |
| *k*<sub>8</sub> = | 0.140625 | *µ*M<sup>−1</sup>s<sup>−1</sup> | EC<sub>50,Cdc25</sub> = | 35 | nM |
| *k*<sub>9</sub> = | 0.25 | *µ*M<sup>−2</sup> | *a*<sub>Wee1</sub> = | 0.4/60 | s<sup>−1</sup> |
| *k*<sub>10</sub> = | 0.025 | s<sup>−1</sup> | *b*<sub>Wee1</sub> = | 2/60 | s<sup>−1</sup> |
| *D*<sub>RT</sub> = | 0.08 | *µ*m<sup>2</sup>s<sup>−1</sup> | *n*<sub>Wee1</sub> = | 3.5 | – |
| *D*<sub>RD</sub> = | 0.4 | *µ*m<sup>2</sup>s<sup>−1</sup> | EC<sub>50,Wee1</sub> = | 30 | nM |
| *D*<sub>F</sub> = | 0.001 | *µ*m<sup>2</sup>s<sup>−1</sup> | *D* = | [2.5, 2.5, 1, 4] | *µ*m<sup>2</sup>s<sup>−1</sup> |
| *α* = | *α*<sub>0</sub> − *s* (*a* − *a*<sub>0</sub>) | – | *γ* = | [2, 7, 4.5, 4.5] | – |
| *α*<sub>0</sub> = | 1 | – | *a*<sub>0</sub> = | 15 | nM |
| *β* = | 0.3 | – | *R*<sub>Nucleus</sub> = | 30 | *µ*m |
| *s* = | 1/50 | nM<sup>−1</sup> |  |  |  |

The geometric and initial-condition controls varied throughout the study are summarized in Fig. 1A and Supplementary Fig. S1. Active and total Cyclin B–Cdk1 are initialized at the same ratio everywhere in the cell, so that the relative balance between Wee1 and Cdc25 activity is spatially uniform. In reality this ratio is unlikely to be uniform, and both the concentration and localization of Cyclin B–Cdk1 change dynamically over the cell cycle ***Gavet and Pines (2010)***; ***Rombouts and Gelens (2021)***; ***Piñeros et al. (2025)***; we keep it fixed for simplicity. The total Cyclin B–Cdk1 concentration in the nucleus is then multiplied by a factor *γ*, the nuclear enrichment factor. Because the activeto-total ratio is held fixed, this raises the nuclear active concentration by the same factor, so that elevated nuclear Cyclin B–Cdk1 seeds the wave while the surrounding cytoplasm starts from the lower baseline. We additionally vary the nuclear radius *R*_Nucleus_, the nuclear position *z_N_* along the animal–vegetal axis, and the diffusion coefficient *D*, which for fixed reaction kinetics is equivalent to changing the effective cell size.

### Cortical Rho–actin

For the cortex we build on an established model of the Rho GTPase network ***Michaud et al. (2022)***, which reproduces a wide range of cortical behaviours (see Box 2D),

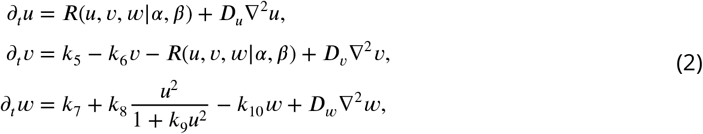

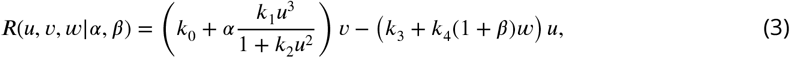

where *u*, *v*, and *w* denote Rho-GTP, Rho-GDP, and F-actin (see Box 2A–C). The parameter *α* represents the effective Ect2 concentration and sets the distance to a subcritical Hopf bifurcation of the spatially homogeneous system; *β* is fixed (Tab. 1). All remaining parameters follow ***Michaud et al. (2022)***.

### Coupling

The two modules are coupled through Ect2, as sketched in Fig. 1A: active cytoplasmic Cdk1 phosphorylates Ect2, reducing its membrane affinity and thereby lowering the effective concentration *α* ***Hara et al. (2006)***; ***Niiya et al. (2006)***; ***Su et al. (2011)***,

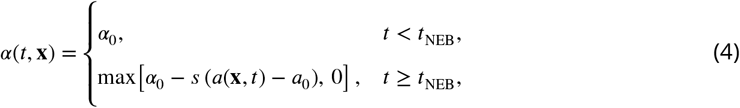

where *s* is the inhibition strength, *a*_0_ the basal active Cyclin B–Cdk1 concentration, and *a*(**x**, *t*) the active Cyclin B–Cdk1 concentration in the cytoplasm from Eq. (1); *α* is bounded below by zero to avoid unphysical negative Ect2 concentrations. The coupling is one-way: we do not include mechanical feedback from the cortex onto the cytoplasm (see Discussion). For comparison we also use an idealized Heaviside inhibition (Methods and Materials, Eq. (8)), which crosses the Hopf bifurcation abruptly rather than inheriting the slow recovery and rapid decay of the Cdk1 waveform.

Having defined the cytoplasmic oscillator, the cortical network, and their coupling, we now turn to the geometry-dependent wave behavior they produce.

## Results

### Directionality of cytoplasmic Cdk1 waves

During both mitosis and meiosis, Cyclin B–Cdk1 accumulates in the nucleus prior to NEB. Upon NEB, active Cyclin B–Cdk1 is released into the cytoplasm, where it propagates spatially and temporally. Capturing this faithfully would require dynamically relocating many cell cycle regulators in space and time, such as Cyclin B, Wee1, and Cdc25, which quickly becomes very complex ***Rombouts and Gelens (2021)***; ***Piñeros et al. (2025)***. We therefore make a deliberate simplification: the same oscillator and the same active-to-total distribution are used throughout the cell, and the nucleus differs only through the activation factor *γ*, which raises the nuclear Cyclin B–Cdk1 concentration at NEB (see Model). This cytoplasmic signal provides the upstream input for the cortical dynamics analyzed in the following section; here we first characterize the cytoplasmic Cdk1 waves themselves.

### Wave fronts, wave backs, and directionality

Following NEB, the spatial Cdk1 profile exhibits a characteristic wave-like structure consisting of two distinct components: an activation front, marking the transition from low to high Cdk1 activity, and a wave back, corresponding to the decay of Cdk1 activity after the peak. Throughout this section, we quantify propagation by tracking the positions of the front and back in kymographs extracted along the radial coordinate *z* from the nucleus toward the cortex (Fig. 1).

From these trajectories, we define the front velocity *v*_F_ and back velocity *v*_B_ as the slopes of the corresponding features in space–time plots (see Supplementary Fig. S2). Positive velocities correspond to propagation away from the nucleus. While *v*_F_ is always positive, *v*_B_ may be either positive or negative, providing a quantitative measure of wave reversibility and defining the sign of back propagation used throughout this section.

To investigate how the sign of *v*_B_ is set, we vary three controls introduced in the model section (Fig. 1A): the nuclear enrichment factor *γ*, nuclear size, and the diffusion coefficient *D*. Because active and total Cyclin B–Cdk1 are initialized at the same ratio everywhere, increasing the nuclear total concentration by *γ* raises the nuclear active concentration by the same factor, so that the nucleus starts enriched relative to the cytoplasmic baseline (see Supplementary Fig. S1). Increasing nuclear size similarly increases the total amount of active Cyclin B–Cdk1 initially contained in the nucleus. As shown below, these two manipulations can be combined into the total nuclear active Cyclin B–Cdk1 content, *N*_nuc_, which determines their effect on wave-back directionality. The third control is *D*, which, for fixed reaction kinetics, is mathematically equivalent to changing the effective system size.

### Baseline regime: traveling-wave–dominated propagation

We first establish a reference regime in which the cytoplasmic Cdk1 wave exhibits traveling-wave– like behavior. At moderate diffusion coefficient and low nuclear enrichment factor, the activation front propagates with approximately constant velocity, producing a linear trajectory in kymographs (constant-speed propagation) (see Fig. 2A (i)). In this regime the wave back follows the same direction as the front, with *v*_B_ > 0, and the overall wave appears as a single traveling entity.

**Figure 2.**
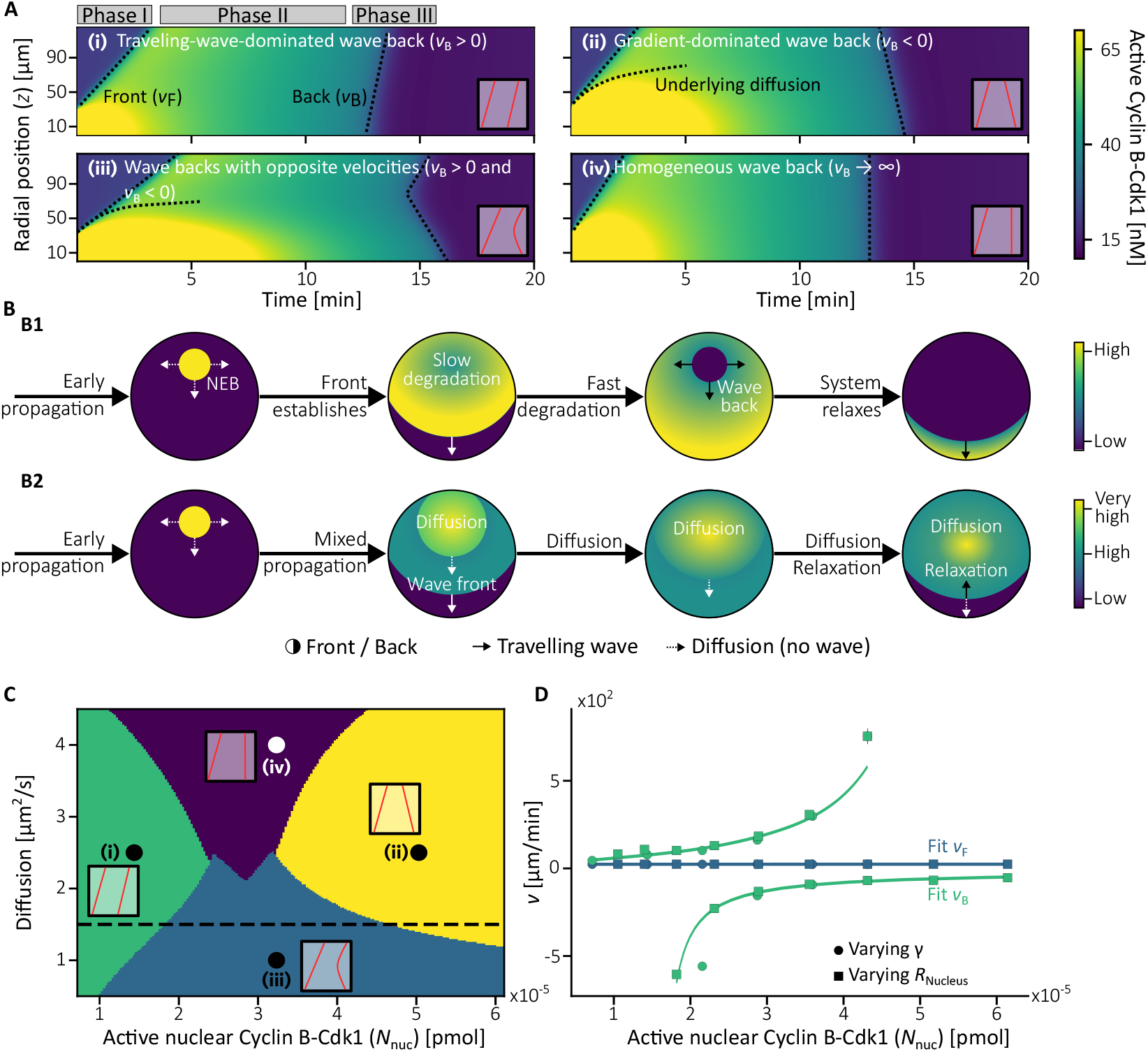
Cyclin B–Cdk1 dynamics: characterization of regimes and wave velocities. **A.** Dynamical regimes observed in the simulations. The wave front remains relatively stable, whereas the wave back behavior changes dramatically. Each regime arises from different initial conditions and diffusion coefficients; the corresponding parameter values are indicated by dots in panel C. **B.** Schematic representation of activity in a generic spatiotemporal system comparing control (*γ* = 1, B1) and increased nuclear enrichment factor (*γ* > 1, B2). Under an increased nuclear enrichment factor, the wave back can reverse direction. **C.** Phase diagram of the distinct dynamical regimes identified in the cytoplasmic system: (i) propagation dominated by traveling waves; (ii) fully reversed wave backs; (iii) coexistence of traveling-wave-dominated fronts with reversed wave backs; and (iv) planar wave backs associated with homogeneous relaxation. Panel D is computed along the black dashed line. **D.** Wave front and wave back velocities as a function of the total active nuclear Cyclin B–Cdk1 content (*N*_nuc_, Eq. 7). This quantity collapses two scenarios onto a single control variable: variation of the nuclear enrichment factor *γ*, and increase of nuclear size (*R*_Nucleus_) at fixed *γ* > 1. Apparent divergences in *v*_B_ arise from vanishing spatial domains over which the velocity is measured (see Supplementary Fig. S3C). Wave back velocities are fitted with a homographic function and the wave front velocities were fitted to a linear function with a quasi-constant velocity of ≈ 23 *μ*m∕min.

### Front–back asymmetry: traveling fronts and gradient-controlled backs

Across a broad range of parameters, we observe a qualitative asymmetry between the wave front and the wave back. While the activation front propagates with nearly constant velocity, consistent with trigger-wave–like behavior, the wave back does not generically exhibit constant-speed propagation. Instead, back trajectories frequently display curvature or piecewise-linear behavior, indicating gradient-controlled dynamics rather than local bistable kinetics (see Fig. 2A (ii), (iii), (iv) and Supplementary Fig. S4A). As a result, the front and back velocities respond differently to changes in nuclear Cdk1 content and system size, allowing the wave back to slow down, stall, or reverse direction even while the activation front continues to propagate outward. This front–back asymmetry is the key mechanism underlying the direction reversals described below.

Although our results do not show substantial changes in the wave front, in the small-system limit the front is expected to become diffusion-dominated as well. The distance beyond which a traveling wave overtakes diffusive spreading can be estimated, for an initially localized region of high concentration diffusing in space, as

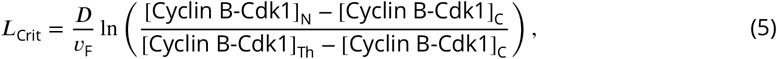

where [Cyclin B-Cdk1]_N_ and [Cyclin B-Cdk1]_C_ are the initial concentrations in the nucleus and in the rest of the cell, and [Cyclin B-Cdk1]_Th_ is the threshold concentration required to excite the system (see Methods and Materials). This diffusion-dominated limit appears as an initial transient even in large systems, but when the system size is much larger than *L*_Crit_ its effect on the measured front velocity is minimal. Consistent with this, we observe an initial diffusion-dominated phase in Supplementary Fig. S2 before a constant-speed front is established. Accordingly, throughout the remainder of the paper we focus on regimes in which a well-defined traveling front exists.

### Nuclear Cyclin B–Cdk1 content controls wave-back directionality

We tested the effect of nuclear Cyclin B–Cdk1 by varying the nuclear enrichment factor *γ*. Keeping the diffusion coefficient fixed at 1.5 *μ*m^2^/s, we found that increasing the nuclear concentration drove the wave back from positive velocity, through a regime of coexisting wave behaviors, to negative velocity (see Fig. 2D). Increasing the nuclear Cyclin B–Cdk1 either by raising its concentration (*γ*) or by enlarging the nuclear radius produced equivalent effects (see Supplementary Fig. S3A), so the two can be collapsed onto the single control variable *N*_nuc_, the total nuclear content of active Cyclin B–Cdk1 (Eq. 7; Fig. 2D and Supplementary Fig. S3A-C). In the transient regime of coexisting velocities, the positive wave-back velocity appears to diverge while the negative velocity converges from large negative values toward the wave-front velocity (see Fig. 2D); this apparent divergence arises because the spatial domain over which each velocity is measured shrinks to zero (see Supplementary Fig. S3D).

Intuitively, when *N*_nuc_ is low the system supports a single traveling wave. As it increases, two processes coexist: a wave front propagating outward at constant speed, and a slower diffusive spread of the elevated nuclear pool. Because diffusion lags the front, regions farther from the nucleus begin to relax before the diffusive component reaches them. This sets up a gradient that drives relaxation inward, from the most distant regions back toward the nucleus, producing a backward-propagating wave back (see Fig. 2B).

### System size and diffusion modulate gradient effectiveness

Because the proposed mechanism for wave-back reversal relies on diffusion-driven gradients, we used the diffusion coefficient as our second control parameter, which is mathematically equivalent to changing the system size for fixed reaction rates. In small systems (large diffusion coefficients), large nuclear concentrations of active and total Cyclin B–Cdk1 diffuse rapidly throughout the domain, producing a nearly homogeneous decay of Cdk1 activity and eliminating directional wave-back propagation (see Fig. 2A (iv)). In larger systems (small diffusion coefficients), gradients persist over longer distances, allowing the wave back to exhibit hybrid behavior consisting of both traveling-wave–dominated and diffusion-driven components (see Fig. 2A (iii)). The range of total nuclear Cdk1 content over which such hybrid behavior occurs narrows as diffusion increases, consistent with the asymptotic behavior of the fits in Fig. 2D and Supplementary Fig. S4E.

### Phase diagram of Cdk1 waves

Together, these results define a phase diagram of the cytoplasmic Cdk1 wave behavior as a function of the total nuclear active Cyclin B–Cdk1 concentration and the diffusion timescale (or system size). Distinct regimes can be identified based on the sign and spatial structure of the wave-back velocity: (i) a traveling-wave–dominated regime with *v*_B_ > 0, (ii) a counterpropagating-back regime with *v*_B_ < 0, (iii) a hybrid regime with coexistence of both behaviors, and (iv) a fast-diffusion regime in which no well-defined wave propagation occurs.

These regimes provide a simple geometric framework for interpreting organism- and stage-specific differences in observed wave directionality. Returning to the two prototypical cell types introduced in the top and bottom rows of Fig. 1B,C, we find that our results are consistent with the distinct characteristics of these systems. An embryo of *Xenopus laevis* during its first mitosis (approximately 1 mm in diameter) can be considered a very large system in comparison with the size of its nucleous. At the same time, a starfish oocyte in meiosis represents a smaller system (approximately 0.15 mm in diameter) with a much larger germinal vesicle. We now turn to the cortical network and examine how Cdk1 waveforms are transmitted to and converted into Rho–actin dynamics in the excitable cortex.

#### Box 2.

Rho-actin dynamics and cortical waves

**Box 2—figure 1.**
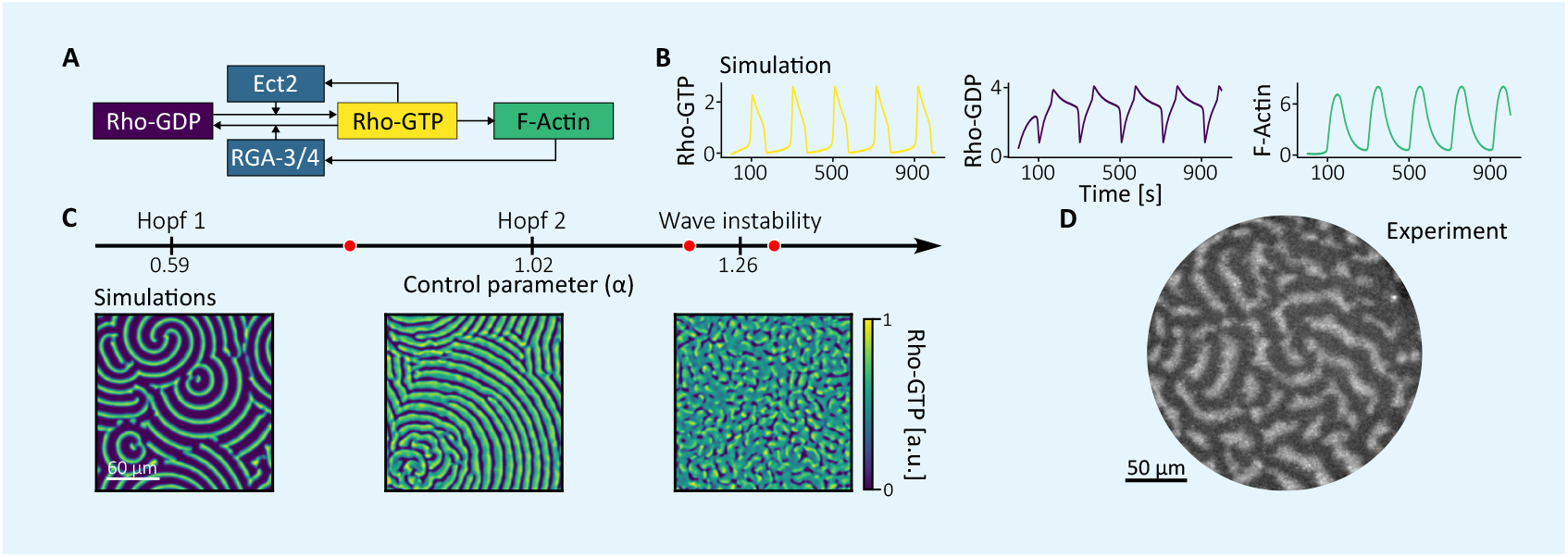
The RhoA network and its dynamics. **A.** Basic network structure regulating RhoA activity. RhoA exists in active, GTP-bound state (Rho-GTP) and inactive, GDP-bound state (Rho-GDP). Activation depends on guanine nucleotide exchange factors such as Ect2, which is itself recruited and activated by Rho–GTP, forming a positive feedback loop ***Rossman et al.*** (2005); ***Goryachev and Leda*** (2019). Inactivation occurs through GTP hydrolysis, a process catalysed by the GTPase-activating protein RGA-3/4 ***Michaux et al.*** (2018). RGA-3/4 is recruited by F-actin, which is polymerised by a formin downstream of Rho–GTP activity, thereby closing a negative feedback loop ***Michaux et al.*** (2018). As in the Cyclin B-Cdk1 system, this combination of positive and negative feedback provides the essential prerequisites for establishing oscillations. **B.** Time series of Rho–GTP, Rho–GDP, and F-actin generated using the model introduced in ***Michaud et al.*** (2022), which is used throughout this paper to reproduce RhoA dynamics. **C.** Bifurcation diagram as a function of the parameter *α*, representing Ect2 concentration and used as a control parameter throughout this paper. Representative spatial patterns of Rho-GTP are shown for *α* = 0.8, 1.2, and 1.3. The model exhibits a rich variety of dynamical regimes, including spirals, wave trains and spiral core turbulence. **D.** Experimental dynamics of active RhoA in a starfish oocytes. Figure adapted from ***Bement et al.*** (2015).

### Cytoplasmic Cdk1 waves shape cortical dynamics

#### The cortex as a Cdk1-gated excitable system

Rather than treating SCWs as autonomous cortical phenomena, we view them as downstream dynamical consequences of cytoplasmic Cdk1 activity. The cortex functions as an excitable system whose dynamics are transiently gated by Cdk1-mediated inhibition. Cytoplasmic Cdk1 is transmitted to the cortex primarily through phosphorylation of the Rho GEF Ect2, which reduces its membrane affinity and suppresses Rho GTPase signaling ***Hara et al. (2006)***; ***Niiya et al. (2006)***; ***Su et al. (2011)***. Changes in Cdk1 activity therefore modulate the distance of the cortical network to a Hopf bifurcation separating quiescent and oscillatory regimes, and SCWs emerge as phase waves—apparent propagation arising from spatially staggered threshold crossing rather than from local signal transmission—as different cortical regions cross this bifurcation at different times ***Wigbers et al. (2021)***. This phase-wave mechanism corresponds to one of the dynamical regimes identified in our previous study of coupled excitable systems with widely separated diffusion timescales ***Cebrián-Lacasa et al. (2024)***. Here, we extend that framework using a biologically realistic model and show that the resulting cortical reactivation can take one of two distinct forms: a planar wave that sweeps across the cortex, or pacemaker-driven recovery that generates target patterns, with circular wavefronts spreading radially outward from localized sites of reactivation. In the remainder of this section, we ask which properties of the cytoplasmic signal and the cortex select between these alternatives.

#### Cortical excitability and control by Ect2

The cortical dynamics are described using an activator–depleted-substrate–inhibitor (ADSI) model of the Rho GTPase network ***Bement et al. (2015)***; ***Michaud et al. (2022)***; ***Chomchai et al. (2025)***. The model tracks Rho-GTP, Rho-GDP, and F-actin; other regulators, including Ect2 and the GAP RGA-3/4, enter through parameters. It captures transient oscillations and a wide range of spatial patterns, with the effective Ect2 concentration *α* as the key control parameter (see Box 2C). For low *α* the cortex is quiescent; increasing *α* induces oscillations and spatial patterning. Two Hopf bifurcations occur within the relevant range, at *α* = 0.59 and *α* = 1.02, but propagating cortical waves persist across the entire active window 0.59 < *α* < 1.26 up to the wave instability at *α* = 1.26. In the absence of Cyclin B–Cdk1 inhibition, a cortex expressing sufficiently high Ect2 (i.e. within this window) therefore does not spontaneously reset on the mitotic timescale, but remains in the wave-propagating regime.

#### Coupling cytoplasmic Cdk1 to cortical dynamics

The interaction between cytoplasmic Cdk1 and the cortex is implemented by inhibiting Ect2, through a reduction in *α*. We consider two inhibitory signals. The first is an idealized Heaviside inhibition that approximates an abrupt on–off Cdk1 signal. The second is derived directly from the cytoplasmic Cdk1 profile and preserves the slow recovery and rapid drop of the Cdk1 waveform after NEB. Both suppress cortical activity, but they differ in how inhibition is released: the Heaviside signal crosses the Hopf bifurcation abruptly, whereas the Cdk1-derived signal recovers slowly and then drops quickly. As we show below, this difference strongly affects how cortical activity reappears.

Alongside the inhibitory signal, we vary how F-actin disassembly is represented. The disassembly term can be either spatially homogeneous or heterogeneous, with heterogeneity representing additional, unmodeled factors that influence F-actin turnover. Together, the inhibition and disassembly types define a 2 × 2 comparison—abrupt versus waveform-derived inhibition, each combined with homogeneous or heterogeneous actin turnover—which we use to determine which combinations produce planar reactivation waves and which produce pacemaker-driven recovery.

#### Cortical response to Heaviside inhibition in two dimensions

Under Heaviside inhibition, cortical activity is rapidly suppressed and the system approaches a homogeneous steady state. After inhibition is lifted, activity reappears through two distinct mechanisms (Supplementary Videos 6,7; Fig. 3D,E). In heterogeneous systems, parameter inhomogeneities allow an immediate recovery of complex dynamics, requiring only a single reactivation wave before spiral dynamics re-emerge. In homogeneous systems, recovery is delayed and proceeds through several wave cycles before spiral turbulence re-emerges. This difference is reflected in the oscillation period, since spiral wave dynamics in excitable systems are characterized by a shorter effective period ***Cebrián-Lacasa et al. (2024)***. It arises because, as the system approaches the stable state during inhibition, phase differences between neighboring points — set earlier by the pre-inhibition pattern — are smoothed out by diffusion. Upon reactivation, a homogeneous system must first re-amplify residual inhomogeneities, whereas in a heterogeneous system the imposed parameter variations immediately seed pattern formation, eliminating this delay.

**Figure 3.**
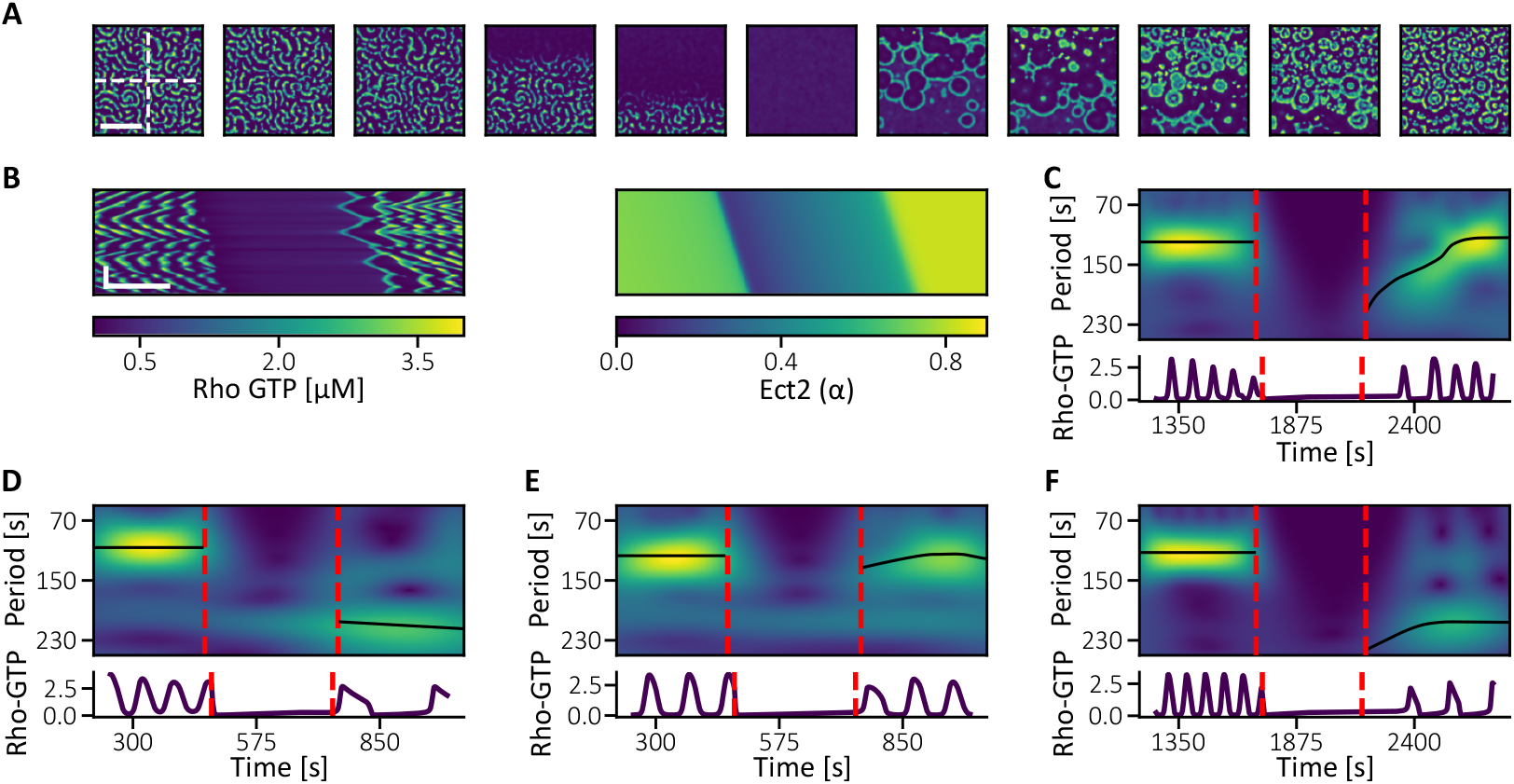
Cortical dynamics and cytoplasmic inhibitory waves. **A.** Frames of Rho-GTP dynamics coupled with a reduction in effective Ect2 caused by Cdk1 inhibition with heterogeneous F-actin disassembly. Scale bar: 50*μ*m. **B.** Kymographs over the vertical dashed line in A of the Rho-GTP concentration and the effective Ect2. Scale bars: 25*μ*m (*y*-axis) and 240s (*x*-axis). **C.** Averaged period over the horizontal dashed line in A using wavelet transformation and time series corresponding to (*x*, *y*) = (150, 150). **D.** Averaged period with homogeneous F-actin disassembly and Heaviside inhibition (Supplementary Video 6). **E.** Averaged period with heterogenous F-actin disassembly and Heaviside inhibition (Supplementary Video 7). **F.** Averaged period with homogeneous F-actin disassembly and Cdk1 inhibition (Supplementary Video 8). Exceptionally *s* = 1∕75*μ*M^−1^

#### Cortical response to Cdk1-derived inhibition in two dimensions

When inhibition follows the cytoplasmic Cdk1 profile, cortical activity is again rapidly driven toward a homogeneous steady state. In a homogeneous system, the dynamics qualitatively resemble the Heaviside case: several planar waves propagate through the system, yielding a longer oscillation period than in the spiral regime (Supplementary Video 8; Fig. 3F). The same mechanism described above therefore applies.

In heterogeneous systems, the behavior is qualitatively different (Supplementary Video 5). As shown in Fig. 3A, local pacemakers generate target patterns that collectively form a propagating reactivation front, rather than a sequence of planar waves. The oscillation period is then set by the target patterns and is comparable to both the planar-wave cycle period and the intrinsic period of the system, rather than to the shorter spiral-wave period (see Fig. 3B,C). After a few oscillation periods, the pattern evolves into a multi-spiral regime, recovering dynamics similar to those observed before inhibition and in the corresponding Heaviside case. As we show next, these differences arise from the combined effects of slow recovery, parameter heterogeneity, and the manner in which the system crosses the Hopf bifurcation.

#### Cortical dynamics in a spherical geometry

We next extend the cortex to a three-dimensional shell and use the cytoplasmic regimes of Fig. 2D as input signals regulating Ect2 (see Fig. 4A; Supplementary Videos 9–12). The resulting cortical Rho GTPase signal reproduces the four cytoplasmic behaviors: a wave back propagating parallel to the front, a counter-propagating wave back, coexistence of the two, and a flat recovery occurring simultaneously across the cell. The first two correspond to the scenarios seen experimentally (Fig. 1D), while the flat recovery may describe surface contraction waves once the embryo has divided several times and cell size is strongly reduced. Even so, the reactivation mode in these simulations does not fully match the experiments of ***Bement et al. (2015)***, where cortical activity reappears after Cdk1 inhibition as an almost planar front (see Fig. 4E; Supplementary Video 18). This discrepancy suggests that the reactivation mode is controlled not only by the spatial ordering of inhibition but also by its temporal profile and strength — which we address next.

**Figure 4.**
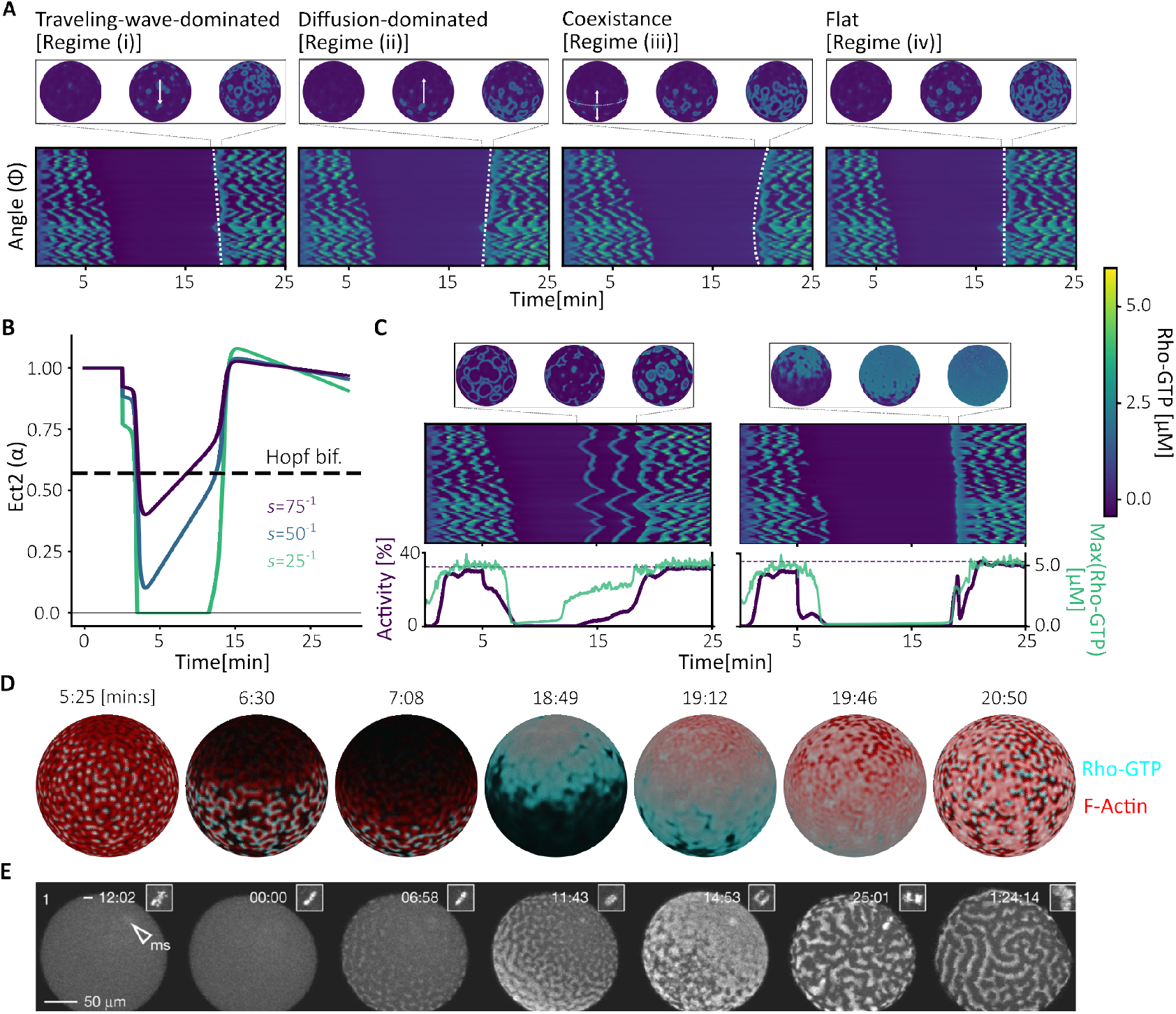
Reconstitution of cortical dynamics during surface contraction waves in a three-dimensional cell cortex. **A.** Cortical dynamic for each of the regimes introduced in Fig. 2. Upon release from inhibition, cortical activity reappears as a moving front whose direction is set by the underlying cytoplasmic Cdk1 waveform (see Supplementary Videos 9-12). **B.** Ect2 signal for different Cdk1 phosphorylation rates (inhibition strengths *s*). **C.** Spatiotemporal dynamics of the corresponding strong and weak inhibition cases given in B. The bottom shows the maximum activity of Rho GTP (green) and the percentage of the system that is active (blue) averaged over the equator of the system. **D.** Frames of a movie with *γ* = 1 showing the merged normalised concentration of Rho GTP (cyan) and F-Actin (red) (see Supplementary Video 20). **E.** Rho GTP signal after Cdk1 inhibition in starfish oocytes. Adapted from ***Bement et al. (2015)***. Full movie in Supplementary Video 18.

#### Effects of inhibition duration and cortical heterogeneity

We therefore returned to the two-dimensional results of Fig. 3. There, Heaviside inhibition could produce planar reactivation waves; however, when F-actin disassembly is homogeneous and inhibition times are long, several wave cycles emerge even in three dimensions, again unlike the experiments (see Supplementary Fig. S6A,B; Supplementary Videos 13,14). Introducing heterogeneous F-actin disassembly instead reproduces a single reactivation wave followed by immediate spiral dynamics (see Supplementary Fig. S6C; Supplementary Video 15). Thus long inhibition in homogeneous systems favors multi-cycle recovery, whereas heterogeneity promotes rapid singlewave reactivation.

#### Inhibition strength selects the reactivation mode

These results suggest that the abrupt release imposed by the idealized Heaviside signal can bias the reactivation mode relative to Cdk1-derived inhibition, under which Ect2 recovers first slowly and then rapidly. To determine how these two recovery phases affect cortical reactivation, we varied the phosphorylation rate, or inhibition strength *s*, with which Cdk1 acts on Ect2 (Fig. 4B,C). Strong inhibition drives the system across the Hopf bifurcation during the fast recovery phase of Ect2, producing an immediate planar phase wave followed by spiral turbulence, in agreement with experiment (Fig. 4C, right, and D; Supplementary Video 17). Weaker inhibition allows the system to cross the bifurcation during the slow recovery phase, so that spatial heterogeneities trigger oscillations locally while other regions remain excitable. This produces target patterns and transient oscillations before the full pattern is re-established (Fig. 4C, left; Supplementary Video 16). This reactivation regime is absent from idealized inhibition models.

#### Reactivation mode shapes the amplitude, extent, and frequency of cortical activity

We next asked whether the two reactivation modes differ in properties that may be relevant to mechanical output. Along the cell equator, we measured the maximum Rho-GTP activity and the active fraction, *f*_act_, defined as the fraction of positions at which Rho-GTP activity exceeds a threshold *u*_th_ = 1 (Fig. 4C). Although target patterns appear earlier during reactivation, they have lower maximum Rho-GTP activity and occupy a smaller fraction of the cortex than the subsequent spiral pattern. Once spiral turbulence is established, both the maximum activity and the active fraction increase. Spiral dynamics also produce more frequent oscillations, corresponding to a shorter effective period than that of target patterns ***Cebrián-Lacasa et al. (2024)***. Thus, the transition from target patterns to spiral turbulence increases the amplitude, spatial extent, and frequency of cortical Rho-GTP activity.

Previous modeling showed that increasing Ect2 can drive the cortical Rho–actin system from an excitable regime characterized by traveling waves into a stable high-Rho state associated with cytokinetic furrow formation ***Goryachev et al. (2016)***. Together with our results, this suggests that the appearance of cortical activity alone may not be sufficient to generate a substantial mechanical response. We hypothesize that the more spatially extensive, higher-amplitude, and higher-frequency activity produced after spiral re-establishment may be more effective at driving cortical contraction. Because our model does not include mechanical coupling, this hypothesis remains to be tested.

#### Alternative ways of controlling Rho GTPase activation

Finally, inhibitory profiles and cytoplasmic activity levels are not the only factors controlling cortical dynamics. Because activity propagates as a concentric wave from the nucleus ***Chang and Ferrell Jr (2013)***, the nuclear position alone can alter both the wave back and the wave front. This is illustrated in Supplementary Fig. S7: moving the nucleus produces a range of behaviors, from a well-defined phase wave when the nucleus lies close to the cortex, to a flat activation profile in which Cdk1 reaches the cortex simultaneously when the nucleus is central (an effectively infinite apparent phase velocity).

## Discussion

Surface contraction waves show how large cells coordinate cortical mechanics with cell cycle progression. These waves travel in opposite directions in starfish oocytes ***Bischof et al. (2017)*** and *Xenopus* embryos ***Chang and Ferrell Jr (2013)***, even though both systems share the same core biochemical machinery. Our work shows that this difference can arise without organism-specific cortical mechanisms. Instead, directionality depends on total nuclear Cdk1 content—which is set jointly by nuclear size and nuclear Cdk1 concentration—and effective cell size. These factors affect the activation front and wave back differently.

Activation fronts behave as trigger waves driven by local bistable dynamics ***Gelens et al. (2014)***; ***Beta and Kruse (2017)***; ***Deneke and Di Talia (2018)***. For fixed reaction and diffusion parameters, they propagate at an approximately constant velocity. Wave backs behave differently: their propagation is shaped mainly by Cdk1 gradients that form after NEB. Because the front and back respond differently to nuclear Cdk1 content and effective system size, they can move at different speeds or even in opposite directions. A large system with a relatively small nuclear Cdk1 pool favors outward co-propagation, as observed during the first mitosis of *Xenopus*. A proportionally larger nuclear pool can instead drive the wave back inward, as in starfish oocytes.

Wave-back reversal requires more active Cyclin B–Cdk1 in the nucleus than in the surrounding cytoplasm at NEB. This model assumption is consistent with experimental data from diverse systems. Several live-cell studies reported rapid, positive feedback-driven translocation of cytoplasmic Cyclin B–Cdk1 into the nucleus just prior to NEBD ***Gavet and Pines (2010)***; ***Santos et al. (2012)***. This positive feedback loop is driven by the activity of the nuclear Cyclin B–Cdk1 complex that promotes translocation of the cytosolic Cyclin B–Cdk1 into the nucleus, where growing nuclear concentration of Cyclin B–Cdk1 further increases its phosphorylation and activation. Gavet and Pines (2010) reported an approximately two-fold nuclear enrichment of Cyclin B1–Cdk1 concentration during this process ***Gavet and Pines (2010)***. Recent experiments in fission yeast further demonstrated that the Cyclin B–Cdk1 orthologue Cdc13–Cdc2 is activated first in the nucleus, where the regulatory network exhibits a stronger bistable response than in the cytoplasm. Subsequent nuclear export of Cdc13–Cdc2 contributes to cytoplasmic Cdk1 activation ***Kapadia and Nurse (2025)***. In frog egg extracts, active Cyclin B–Cdk1 accumulates in the nucleus before activation of the surrounding cytoplasm ***Maryu and Yang (2022)***; ***Puls et al. (2024)***; ***Piñeros et al. (2025)***, and nuclei can determine where mitotic waves begin ***Afanzar et al. (2020)***; ***Nolet et al. (2020)***. Cyclin B enrichment near the nuclear region has also been observed during starfish oocyte meiosis ***Terasaki et al. (2003)***. Further experimental work is necessary to quantify the exact nuclear-to-cytoplasmic difference in active Cyclin B–Cdk1 concentration and establish whether it is sufficient for complete wave-back reversal.

At the cortex, Cdk1-dependent phosphorylation of Ect2 links the cell cycle oscillator to the Rho– actin network by reducing Ect2 membrane affinity ***Hara et al. (2006)***; ***Niiya et al. (2006)***; ***Su et al. (2011)***. As this inhibition is released, different cortical regions cross a Hopf bifurcation at times set by the cytoplasmic Cdk1 waveform ***Wigbers et al. (2021)***. In this regime, the large-scale reactivation front does not need to spread through successive local excitation along the cortex. Instead, the apparent movement results from different regions reactivating one after another. The cortical response can therefore be a phase wave whose direction is inherited from the cytoplasmic signal, consistent with the mechanism proposed in our previous work ***Cebrián-Lacasa et al. (2024)***.

The timing of inhibition release also determines the form of cortical reactivation. Rapid release causes neighboring regions behind the advancing Cdk1 wave back to cross the bifurcation within a short interval, producing a planar reactivation wave consistent with observations in starfish oocytes ***Bement et al. (2015)***; ***Liu et al. (2025)***. When release is slower, spatial differences in actin turnover become important. Some regions reactivate earlier and act as pacemakers, producing transient target patterns before spiral dynamics return. This pacemaker-driven regime does not appear with idealized on–off inhibition in our simulations. Inhibition strength determines when reactivation occurs: strong inhibition keeps the cortex quiescent until the fast recovery phase, whereas weaker inhibition allows heterogeneous regions to reactivate during the slow phase. Together, inhibition strength, release kinetics, and cortical heterogeneity control the reactivation pattern. The broader, stronger, and more frequent activity of the later spiral regime may also be more effective at generating cortical contraction, although this remains a hypothesis because the model does not include mechanics.

Mechanical feedback may still modify these dynamics. Cortical tension, cytoplasmic flows, and surface deformation can influence SCW propagation ***Miller et al. (2018)***; ***Klughammer et al. (2018)***; ***Yin et al. (2022)***, and mechano-chemical coupling contributes more broadly to actin-wave dynamics ***Beta et al. (2023)***. Our results show that biochemical signaling alone can generate the main behaviors in the model, but they do not show that mechanical feedback is unnecessary in living cells. This distinction could be tested by perturbing actomyosin contractility while monitoring Cdk1 and Rho-GTP dynamics. One could then ask whether wave directionality and reactivation mode still follow the biochemical predictions when cell deformation is altered.

Taken together, our results identify two sets of controls. Total nuclear Cdk1 content and effective cell size control cytoplasmic wave directionality, whereas inhibition kinetics and cortical heterogeneity control cortical reactivation. Increasing nuclear active Cyclin B–Cdk1 content should slow, stall, and eventually reverse the wave back, while rapid diffusion should favor spatially uniform recovery. At the cortex, rapid release or strong inhibition should favor a single planar reactivation wave. Slow release combined with cortical heterogeneity should favor pacemaker-driven target patterns, whereas prolonged inhibition in a homogeneous cortex should favor multi-cycle planar recovery. Similar front–back decoupling may occur in other excitable or oscillatory systems in which a localized source generates an activation front while diffusion-driven gradients control recovery. This may represent a broader principle for spatial coordination inside cells.

## Methods and Materials

### The cytoplasmic Cyclin B–Cdk1 model

We model cytoplasmic Cyclin B–Cdk1 dynamics with the spatially extended oscillator of ***Yang and Ferrell Jr (2013)***; ***Puls et al. (2024)***, given in Eq. (1) of the main text, which tracks the total and active concentrations *c*(**x**, *t*) and *a*(**x**, *t*) with feedback through Cdc25, Wee1, and APC/C. The diffusion coefficient *D* is taken equal for the active and total forms for simplicity; allowing distinct coefficients does not qualitatively change the results (data not shown). The ultrasensitive regulatory terms are the Hill functions

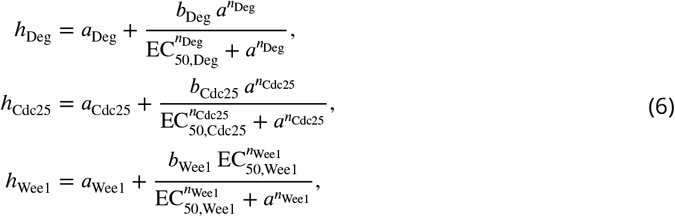

with all parameters taken from ***Yang and Ferrell Jr (2013)***, within experimentally measured ranges ***Tsai et al. (2014)*** (Table 1). The model exhibits relaxation oscillations: rapid activation of Cyclin B–Cdk1 followed by slower, APC/C-driven decay.

To model wave propagation following nuclear envelope breakdown (NEB), Eq. (1) was solved on a one-dimensional radial domain while retaining the three-dimensional diffusive Laplacian under the assumption of spherical symmetry. The origin was placed at the center of the nucleus, and no-flux (Neumann) boundary conditions were imposed at the nuclear boundary. To prevent artificial boundary effects when projecting the cytoplasmic activity onto the asymmetric cortex, the simulation was solved over an extended radial domain. This ensures that all regions of the projected cortex remain unaffected by the outer simulation boundary.

Initial conditions were chosen with active and total Cyclin B–Cdk1 in the same ratio everywhere in the cell, so that the Wee1/Cdc25 balance is spatially uniform. The nuclear concentrations were then raised above the cytoplasmic baseline by multiplying the initial values within the nuclear region by a constant factor *γ*; since the active-to-total ratio is preserved, this raises the nuclear active concentration by the same factor. This is a modeling simplification: in reality the nuclear-to-cytoplasmic distribution of Cyclin B–Cdk1 changes dynamically through the cell cycle ***Rombouts and Gelens (2021)***; ***Piñeros et al. (2025)***; ***Maryu and Yang (2022)***. The total active nuclear content used as the control variable in the main text,

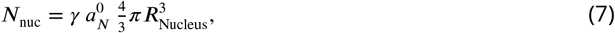

follows directly from this construction.

Wave fronts were defined as the first threshold crossing of active Cyclin B–Cdk1 during activation, and wave backs as the return crossing of the same threshold during decay (the threshold was set to the midpoint between the low and high activity steady states unless otherwise stated). Wave velocities were extracted from space–time kymographs by linear regression over the region where trajectories were approximately linear; in regimes with curved trajectories, piecewise fits were used as described in Supplementary Fig. S2.

System-size effects were explored by varying *D*. Increasing *D* increases the characteristic diffusion length over a fixed timescale and is mathematically equivalent to decreasing system size for fixed reaction kinetics and domain geometry (after nondimensionalization, only the ratio D∕T sets the effective length scale).

### The cortical ADSI model and bifurcation structure

Cortical dynamics were described using the activator–depleted-substrate–inhibitor (ADSI) model of ***Michaud et al. (2022)***, given in Eqs. (3)–(4) of the main text, for active Rho GTPase (*u*), inactive Rho GTPase (*v*), and F-actin (*w*). The three species retain their distinct diffusion coefficients *D_u_*, *D_v_*, *D_w_* (Table 1). The parameter *α* represents the effective Ect2 concentration and sets the distance to a subcritical Hopf bifurcation of the spatially homogeneous system; once this bifurcation is crossed, oscillation amplitudes rapidly increase, giving relaxation-like cortical dynamics that support excitable wave propagation and spiral turbulence. All parameters are retained from ***Michaud et al. (2022)***, within experimentally supported ranges (Table 1).

Spatial heterogeneities in actin turnover were introduced by adding Gaussian-correlated noise to the F-actin degradation rate *k*_10_. The noise field had a fixed correlation length and amplitude (see Numerical methods) and was held constant in time (quenched disorder). These heterogeneities locally shift the Hopf threshold, allowing some regions to enter the oscillatory regime before others, which seeds localized pacemaker regions and target-like domains during cortical reactivation as the system is slowly driven across the bifurcation by Cdk1-dependent inhibition.

### Coupling cytoplasmic Cdk1 to cortical inhibition

Cytoplasmic Cyclin B–Cdk1 was coupled to the cortex as an inhibitory effect on the effective Ect2 concentration *α*, consistent with evidence that Cdk1 phosphorylation reduces Ect2 membrane association ***Hara et al. (2006)***; ***Niiya et al. (2006)***; ***Su et al. (2011)***: Cdk1 activity transiently shifts the cortex across the Hopf bifurcation separating quiescent and oscillatory regimes. The waveform-derived inhibition is given by Eq. (4) of the main text, with *α* bounded below by zero to avoid unphysical negative Ect2 concentrations.

For comparison, an idealized Heaviside inhibition was also used,

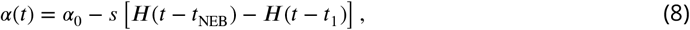

where *t*_1_ − *t*_NEB_ is the inhibition duration. This produces an abrupt crossing of the Hopf bifurcation, in contrast to the waveform-derived inhibition, which yields a slow recovery followed by a rapid decay in *α*(*t*).

In homogeneous cortical systems, longer inhibition drives the system closer to the stable fixed point, reducing residual spatial inhomogeneities; upon release this gives multiple planar traveling waves before spiral turbulence re-emerges, whereas shorter inhibition or a steeper release synchronizes reactivation and favors a single planar phase wave. In systems with heterogeneous actin degradation, localized pacemaker regions persist during inhibition, enabling immediate reactivation regardless of duration; when inhibition is released slowly, these regions cross the Hopf bifurcation before their surroundings and seed target-like patterns. These behaviors are consistent with recovery dynamics in excitable media ***Cebrián-Lacasa et al. (2024)***.

Cortical dynamics were also simulated on a spherical surface representing an early embryonic cell. The cortical domain was a closed spherical shell with no-flux boundary conditions, and cytoplasmic inhibition was applied through the spatiotemporal Cdk1 field projected onto the cortex. Inhibition was driven by the Cdk1 field originating from the off-center nuclear region, positioned along the animal–vegetal axis. Shifting this position altered the relative timing of inhibition across the cortex; in the limit of a centrally located nucleus, all cortical regions are inhibited simultaneously, corresponding to a diverging apparent phase velocity (a spatially uniform phase reset). These effects reproduce the experimentally observed transitions between U-shaped and V-shaped SCWs ***Chang and Ferrell Jr (2013)***.

### Model parameters

In this study, we used two models and retained the parameter values from their original publications ***Yang and Ferrell Jr (2013)***; ***Michaud et al. (2022)***. These values are listed in Table 1 and were used throughout unless explicitly stated otherwise.

In addition to the parameters in Table 1, the spatial parameters are defined as follows. For the one-dimensional cytoplasmic model, the domain radius is set to *R*_Max_ = 125 *μ*m with a nuclear radius of *R*_Nucleus_ = 30 *μ*m. This domain radius is chosen to allow the projection of cytoplasmic activity at any distance onto the cortex. The cortex is defined by the cell radius (*R*_Cell_ = 75 *μ*m), within which the nucleus is positioned 40 *μ*m from the center along the animal–vegetal axis.

Heterogeneities in the cortical model were generated using spatially correlated Gaussian noise applied to the F-actin degradation rate *k*_10_, with a correlation length of 4 *μ*m and a relative amplitude of 2 (i.e. the standard deviation of the noise field, normalized to the baseline *k*_10_; the variance is set by the noise-generation procedure described in the code). The control parameter was held at *α* = *α*_0_ = 1 prior to nuclear envelope breakdown (NEB), which occurred at *t*_NEB_ = 5 min.

### Numerical methods

Both models were integrated in time using an explicit second-order Runge–Kutta method, implemented in custom code. For spatial discretization, the cytoplasmic model and the two-dimensional cortical simulations used a finite-difference scheme, while the three-dimensional cortical simulations used a finite-element method implemented in FEniCSx.

Unless otherwise stated, the spatial and temporal discretization parameters (grid spacing, time step, and domain resolution) were kept fixed across simulations and are provided in the code referenced in the *Code Availability* section.

The cytoplasmic model was solved on a one-dimensional radial domain centered on the nucleus, retaining the three-dimensional diffusive geometry through the radially symmetric Laplacian

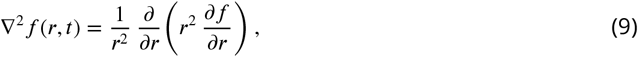

where *f*(*r*, *t*) is radially symmetric about the nucleus. This reduces the dynamics to the radial coordinate while preserving three-dimensional diffusion. Because the nucleus is positioned off-center, the distance from the nucleus to the cortex varies with the angular coordinate *φ*; evaluating the radial solution at the cortical radius therefore yields an angle-dependent cortical input, which is the origin of the *φ*-dependent arrival times shown in Fig. 1D. No-flux (Neumann) boundary conditions were imposed at the nuclear boundary. To prevent artificial boundary effects when projecting the cytoplasmic activity onto the asymmetric cortex, the simulation was solved over an extended radial domain (*R*_Extended_ = 150 *μ*m). This ensures that all regions of the projected cortex remain unaffected by the outer simulation boundary. While this extended boundary also uses a Neumann condition, its influence does not affect the physical domain of interest *R*_Max_ = 125 *μ*m shown in the figures.

### Supplemental material

The critical length beyond which trigger-wave propagation outpaces diffusion can be derived as follows. We approximate the elevated nuclear pool as a point source (valid in the far field, *x* ≫ *R*_Nucleus_) diffusing into a cytoplasmic background *C*_C_, so that the concentration profile is

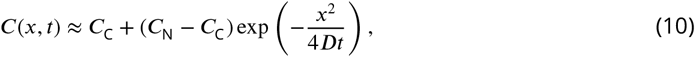

where *C*_N_ is the initial nuclear concentration. Requiring the concentration to reach the excitation threshold *C*_Th_, we solve *C*(*x*, *t*) = *C*_Th_ for the threshold-crossing position,

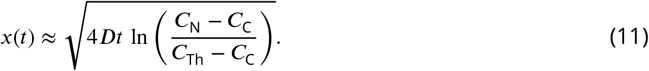

The speed at which this threshold crossing advances follows by differentiation,

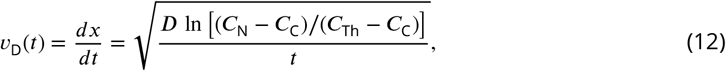

and decreases as *t*^−1∕2^. The trigger wave, by contrast, advances at the constant front velocity *v*_F_. Equating *v*_F_ = *v*_D_ defines the crossover time

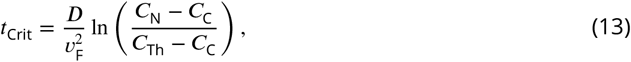

and, using *L*_Crit_ = *v*_F_ *t*_Crit_, the critical length

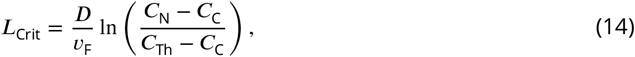

which is the expression quoted in the main text (with *C* ≡ [Cyclin B-Cdk1]). Below *L*_Crit_ the front is diffusion-limited; above it, propagation proceeds at the constant trigger-wave speed *v*_F_.

**Figure S1.**
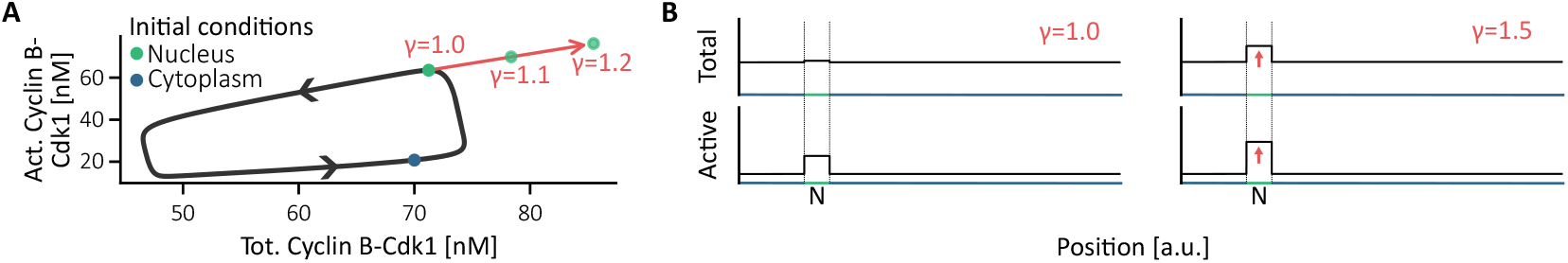
Initial conditions and the nuclear enrichment factor (*γ*). **A.** Limit cycle of the Cyclin B–Cdk1 model with standard parameters, showing the initial conditions in the nucleus and the cytoplasm (green dots). The nuclear enrichment factor *γ* scales the nuclear concentration above the cytoplasmic baseline at fixed active-to-total ratio, as indicated by the red arrow. **B.** Profiles of the initial conditions for *γ* = 1 (baseline) and *γ* = 1.5.

### Supplemental figures and videos

Supplementary Figure S1 illustrates the limit cycle of the temporal Cyclin B–Cdk1 model and the choice of initial conditions, which are selected to lie on the limit cycle with low active Cyclin B–Cdk1 in the cytoplasm and high active Cyclin B–Cdk1 in the nucleus. The factor *γ* then proportionally increases the nuclear concentrations of both total and active Cyclin B–Cdk1 while keeping their ratio, and thus the active fraction, constant.

**Figure S2.**
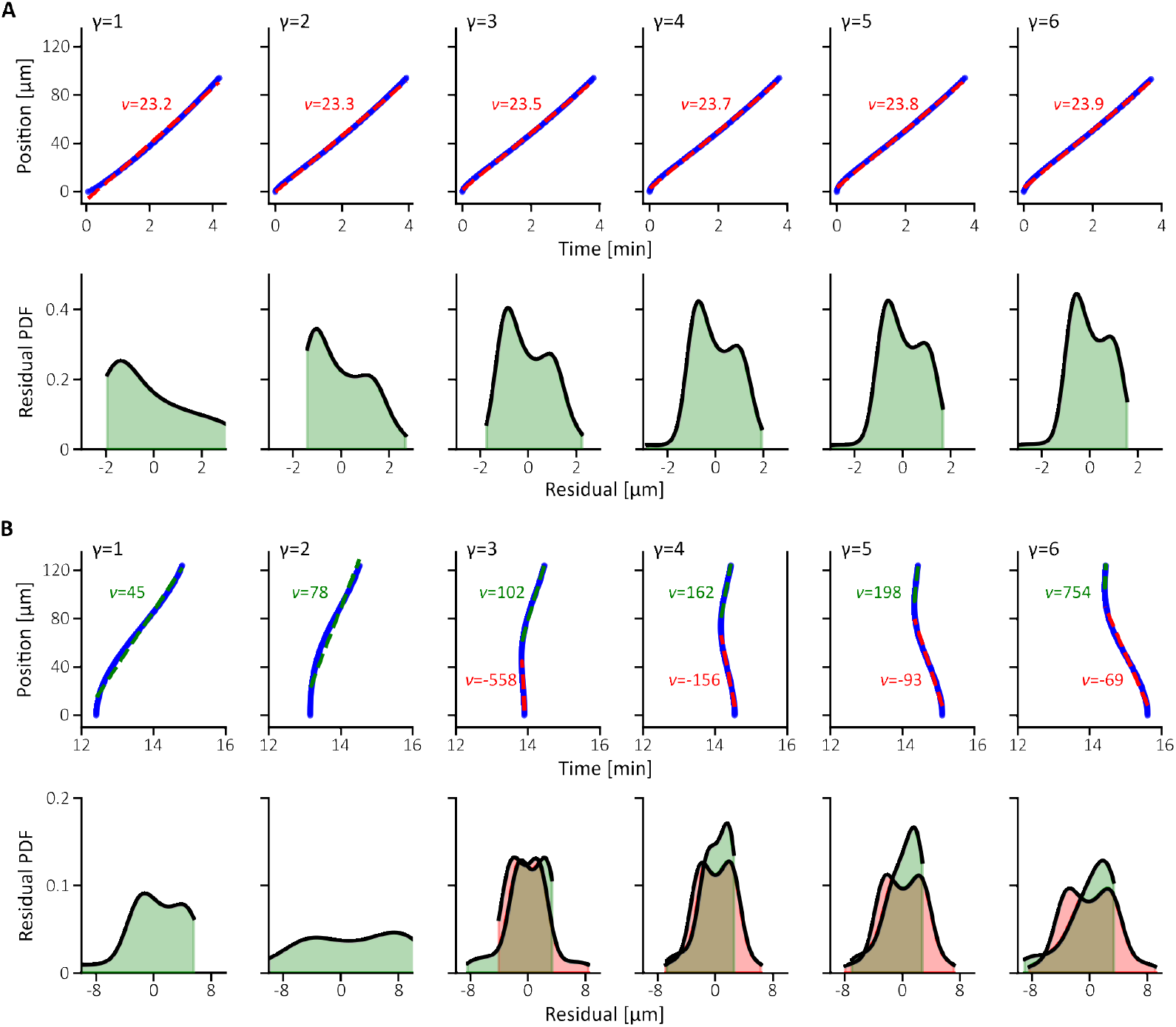
Fits of the wave fronts (A) and wave backs (B) for different nuclear enrichment factors (*γ*) at constant diffusion (*D* = 2.5 *μ*m^2^s^−1^).

Supplementary Figure S2 shows the fits and residuals of the wave fronts and wave backs for different values of the nuclear enrichment factor, corresponding to a subset of the velocities shown in Fig. 2D. The fits were performed using NumPy’s `polyfit` with a first-order polynomial, applied either globally or piecewise (one or two segments) depending on the behavior.

**Figure S3.**
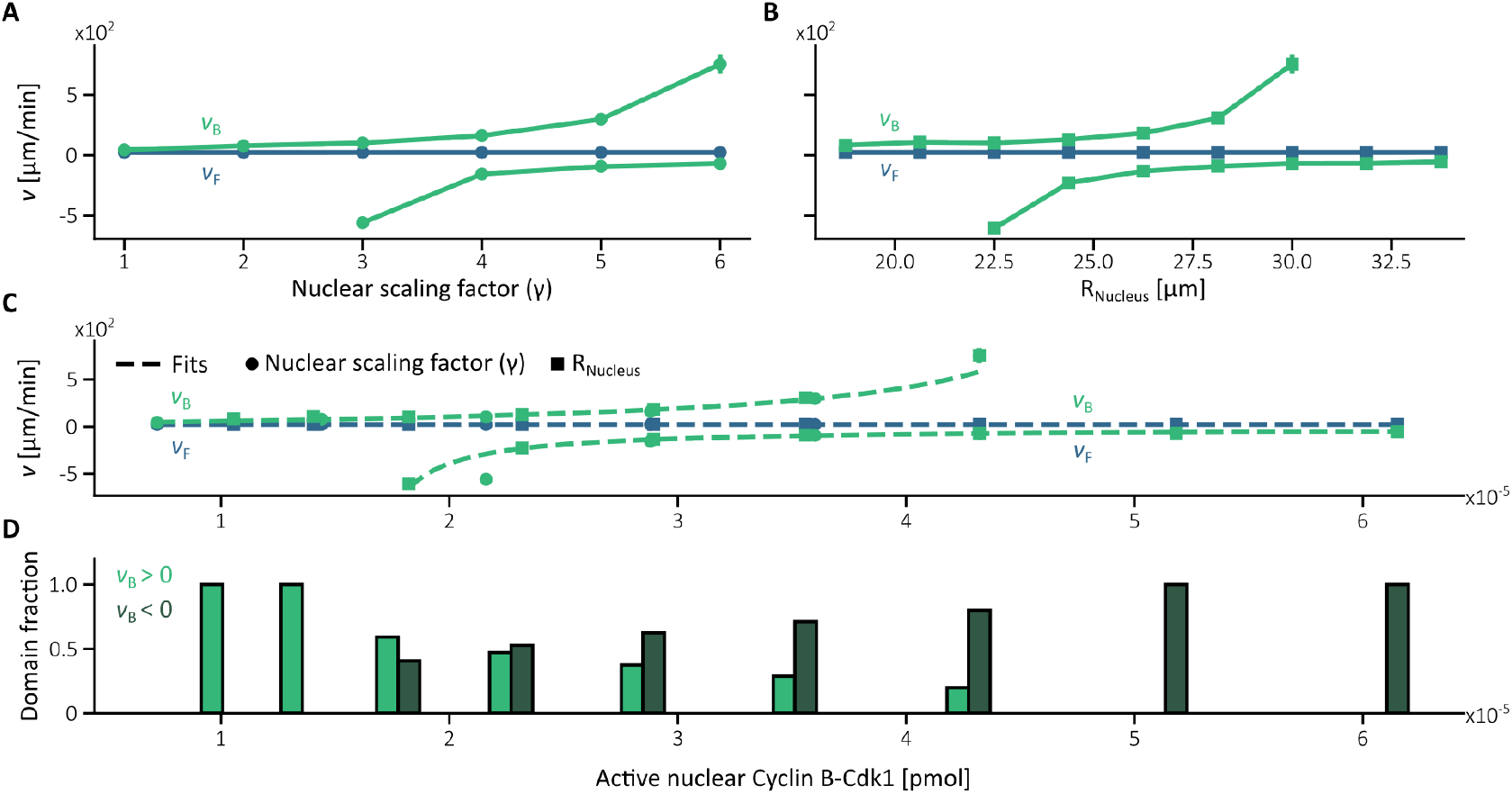
Wave velocity as a function of total active nuclear Cyclin B–Cdk1 (*N*_nuc_) for *D* = 1.5 *μ*m^2^∕s. Wave-front and wave-back velocities for: **A.** varying nuclear enrichment factor (relative nuclear size = 0.24), and **B.** varying relative nuclear size (nuclear enrichment factor = 2). **C.** Data from panels A and B collapsed onto *N*_nuc_. **D.** Fraction of the domain exhibiting traveling-wave–dominated versus diffusion-driven wave-back dynamics.

As expected, the wave fronts exhibit an approximately constant velocity, reflected in good linear fits. However, as *γ* increases, the residuals become bimodal, reflecting an initial phase dominated by diffusion rather than by a traveling front. This also explains the slight variation in fitted velocity, and indicates that the front dynamics decompose into two phases: an initial diffusion-driven segment followed by a traveling-front-driven segment.

In contrast, the wave backs are poorly approximated by a single linear fit, because the established Cdk1 gradient dominates the dynamics. As *γ* increases, the linear approximation becomes less accurate due to the increasingly diffusive character of the gradient, and two segments with opposite apparent velocities can be identified.

We next examined how this hybrid wave back, consisting of traveling-wave–dominated and diffusiondriven segments, varies (Supplementary Fig. S3). We first verified that the transition in wave-back behavior is controlled by the total nuclear Cyclin B–Cdk1 content by modifying the nuclear size rather than the nuclear concentration (via *γ*). Both manipulations produce equivalent behavior after renormalization, collapsing onto the single control variable *N*_nuc_. We then quantified the transition by measuring the fraction of the domain governed by traveling-wave–dominated versus diffusion-driven dynamics.

To complement Fig. 2, Supplementary Fig. S4A–C illustrates the effect of system size—equivalently, the diffusion coefficient—on wave dynamics. A large diffusion coefficient (small effective system) gives a nearly homogeneous distribution of active Cyclin B–Cdk1, so no well-defined wave propagation occurs. A small diffusion coefficient (larger system) produces a spatial gradient confined to part of the domain, giving a wave back with both counterpropagating and forward-propagating segments; as the system size increases further, the forward-propagating component dominates. To characterize these transitions, we quantified wave velocities for different nuclear enrichment factors (Supplementary Fig. S4D) and how the coexistence region of oppositely propagating wave-back segments narrows as diffusion increases. This region is bounded by the two asymptotes of the homographic fit, which converge as the diffusion coefficient increases (Supplementary Fig. S4E).

**Figure S4.**
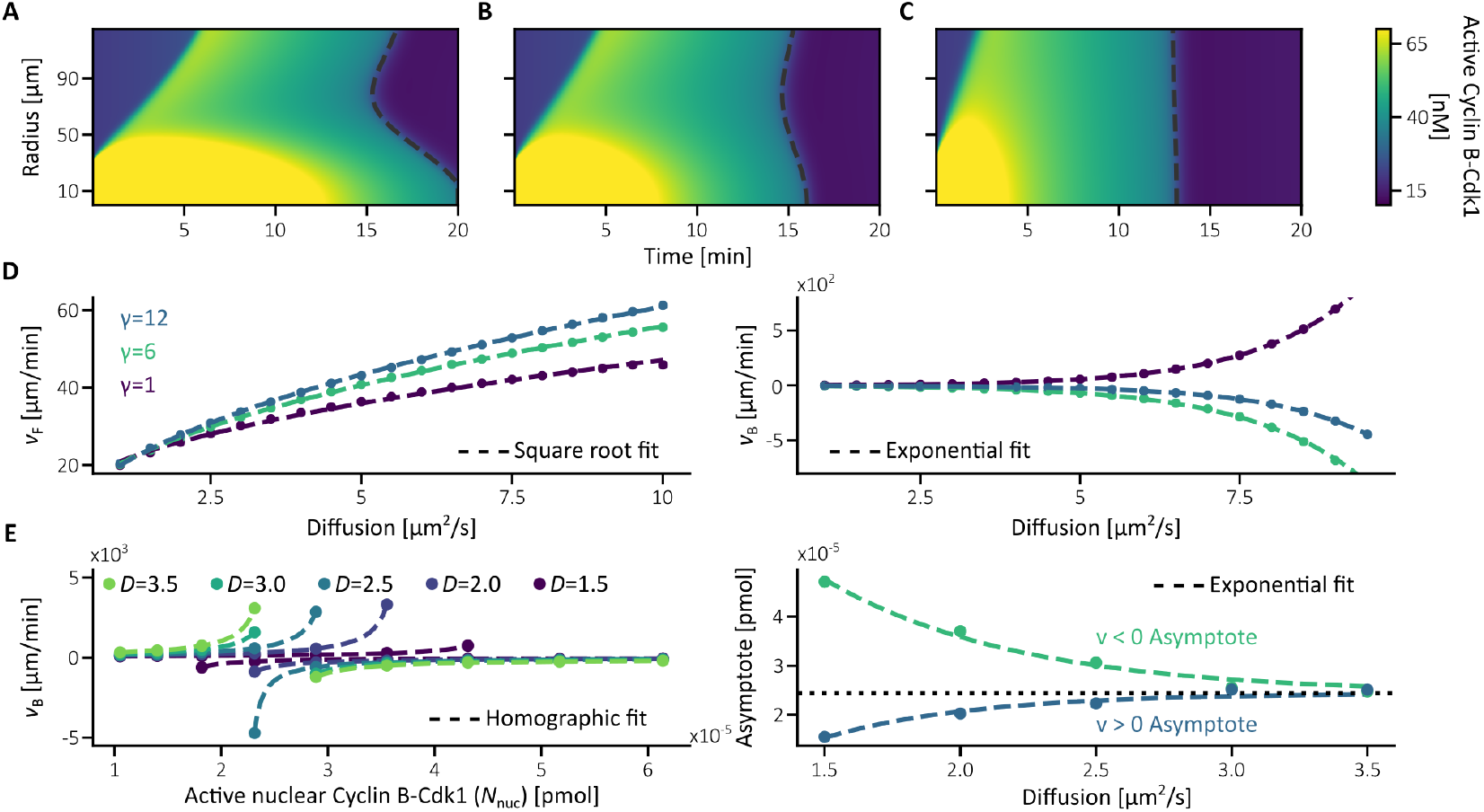
Effect of system size on active Cyclin B–Cdk1 dynamics. Kymographs for: **A.** *D* = 0.5 *μ*m^2^∕s; **B.** *D* = 1.5 *μ*m^2^∕s; **C.** *D* = 4.5 *μ*m^2^∕s. Small systems (high diffusion; panel C) show a nearly homogeneous decay of Cyclin B–Cdk1 activity, because diffusion rapidly smooths spatial gradients. In larger systems (low diffusion; panel A), gradients persist, giving a combination of counterpropagating wave-back segments and forward-propagating traveling waves. Panels A–C use *γ* = 6. **D.** Wave-front and wave-back velocities for different nuclear enrichment factors. Increasing *N*_nuc_ slows the wave back and eventually reverses it, as larger amounts of Cyclin B–Cdk1 must be degraded. **E.** Homographic fits of wave-back velocities for different diffusion coefficients (left). The coexistence region of oppositely propagating wave-back segments narrows as diffusion increases (right); it is bounded by the two asymptotes of the homographic fit, which converge as the diffusion coefficient increases. This panel uses *γ* = 6.

**Figure S5.**
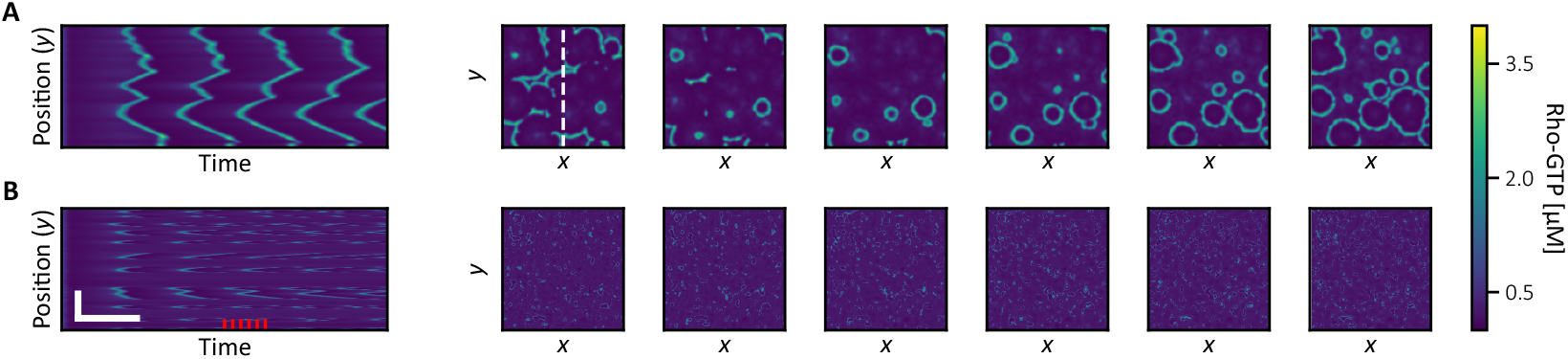
Example of target patterns for *α* = 0.55 with heterogeneous F-actin degradation. **A.** Kymographs (along the white dashed line) and corresponding frames (at the red markers indicated in the kymograph). Regions of higher activity act as pacemakers that emit target-like waves into less active or quiescent regions. **B.** Same simulation as in A but with diffusion set to zero. Local oscillation periods vary across the domain, and some regions do not oscillate, establishing the spatial heterogeneity that seeds target-like domainspattern formation. Scale bars: 25 *μ*m and 180 s.

In Supplementary Fig. S5 we show how operating close to the Hopf bifurcation, in the presence of heterogeneous F-actin degradation, leads to the target-like domains seeded by local pacemakers introduced in the main text. This behavior arises from the excitable nature of the cortical system combined with spatial differences in local oscillation periods (Supplementary Fig. S5B). Regions with shorter intrinsic periods act as pacemakers that excite neighboring regions, generating outward-propagating waves from these localized sources (Supplementary Fig. S5A).

Supplementary Fig. S6 examines the effect of inhibition duration under idealized homogeneous actin degradation. Here spatial patterns emerge only from inhomogeneities in the dynamical variables, since the degradation term is uniform. During inhibition the system relaxes toward its steady state. For short inhibition periods, residual spatial inhomogeneities remain large and pattern reactivation is rapid (Supplementary Fig. S6A). For longer inhibition periods, these inhomogeneities are further reduced by diffusion, and several planar waves appear before spiral or turbulent dynamics re-emerge (Supplementary Fig. S6B). When F-actin degradation is heterogeneous, parameter inhomogeneities alone are sufficient to restart the pattern, giving immediate reactivation regardless of inhibition duration (Supplementary Fig. S6C); the heterogeneities then act as pacemakers that initiate and coordinate reactivation, often before inhibition has fully ended.

**Figure S6.**
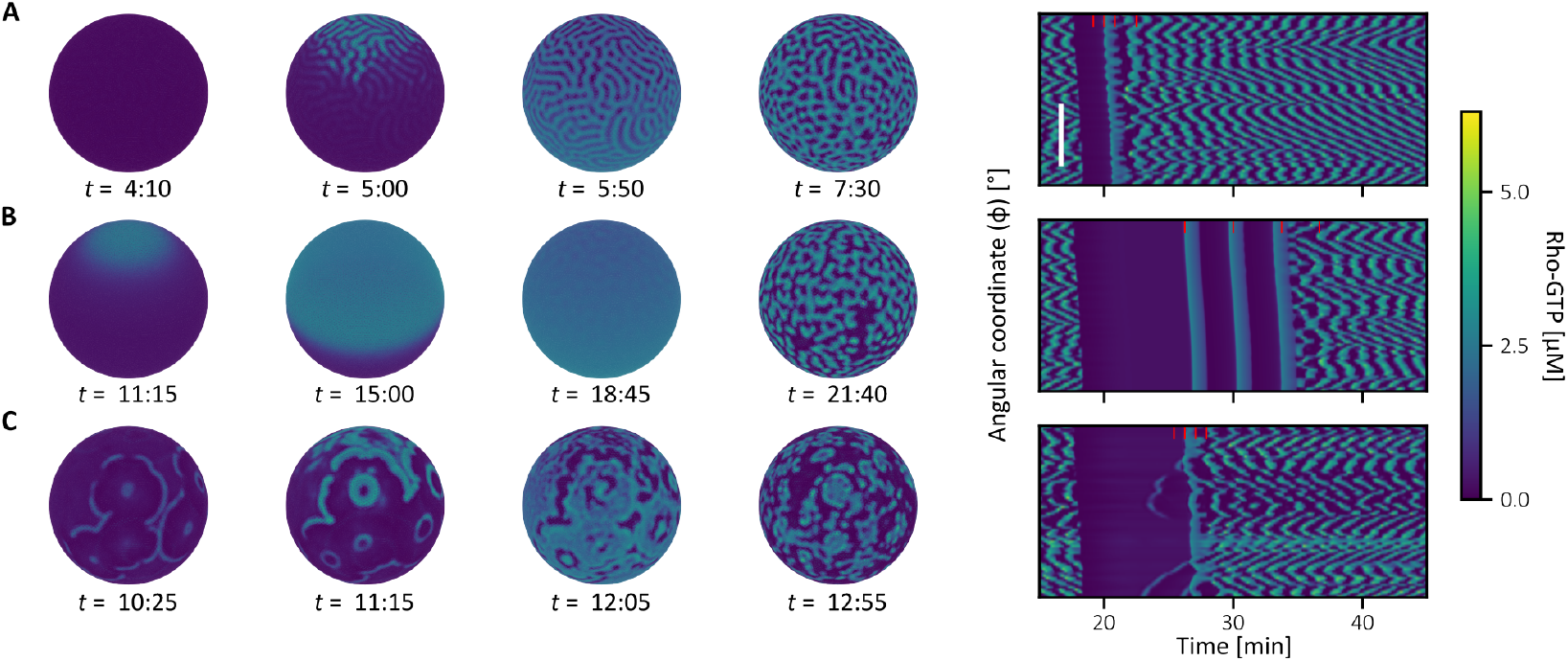
Effect of inhibition duration on the reactivation phase wave. **A.** Short inhibition times preserve spatial inhomogeneities in the dynamical variables, leading to rapid pattern reactivation. Scale bar: 60^◦^ (angular coordinate *φ*, *y*-axis). **B.** Long inhibition times drive the system closer to a homogeneous state; upon release, planar traveling waves appear before heterogeneities are re-amplified and the pattern reactivates. **C.** The previous behaviors occur only when F-actin degradation is homogeneous. When degradation is heterogeneous, parameter inhomogeneities are sufficient to reactivate the pattern immediately. Simulations were performed with *D* = 1.5 *μ*m^2^∕s and *γ* = 1.

**Figure S7.**
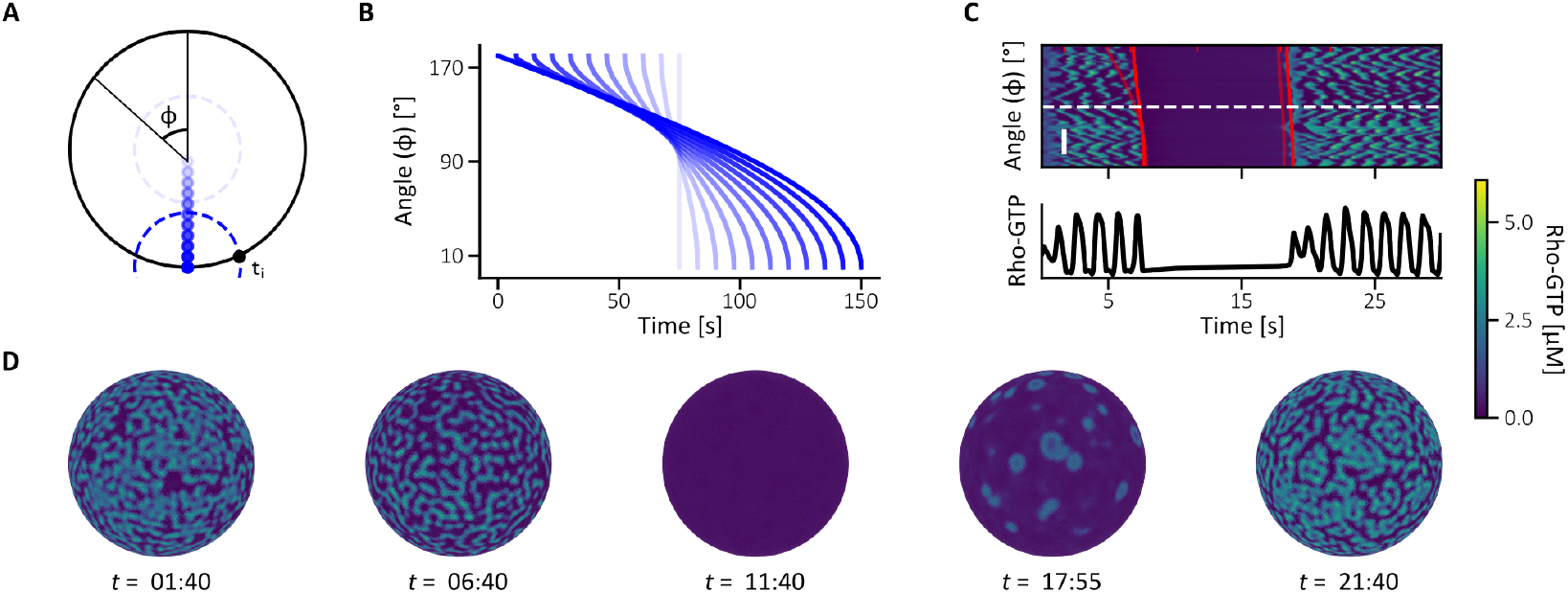
Effect of nuclear position on the reactivation phase wave. **A.** Schematic of the spherical system, with the definition of the coordinates and nuclear positions (rotational symmetry assumed). **B.** Time at which the inhibitory wave reaches each point on the sphere, for different nuclear positions. **C.** Kymograph for *z*_nucleus_ = 7.5 *μ*m (0.10 *r*), with the Ect2 front in red; the semi-transparent curve shows the reference case *z*_nucleus_ = 30 *μ*m (0.40 *r*). Scale bar: 30^◦^ (angular coordinate *φ*, *y*-axis). **D.** Frames corresponding to the kymograph in C. Simulation parameters: *D* = 1.5 *μ*m^2^∕s, *γ* = 1, sphere radius *r* = 75 *μ*m.

The nuclear position directly affects how the wave appears at the embryo surface (Supplementary Fig. S7). In panel C we compare the front and back of the Ect2 wave and show that activation is more synchronous when the nucleus is closer to the center (*z*_nucleus_ = 7.5 *μ*m). The frames in panel D further show that, in this configuration, cortical activity disappears and re-emerges more uniformly than in the off-center case of the main text.

Supplementary videos corresponding to the main and supplementary figures are listed below.

**Video 1.** Projection of Cyclin B–Cdk1 activity onto the surface of a sphere in the traveling-wave–dominated regime (i) (Fig. 2A).

**Video 2.** Projection of Cyclin B–Cdk1 activity onto the surface of a sphere in the front–back asymmetry regime (ii) (Fig. 2A).

**Video 3.** Transition from a labyrinthine to a spiral pattern for *α* = 0.8 (Fig. 3C).

**Video 4.** Target-like domains regime in an overall non-oscillatory system, with parameter heterogeneities acting as pacemakers for *α* = 0.55 (Supplementary Fig. S5).

**Video 5.** Rapid spiral dynamics induced by heterogeneities for *α* = 0.8.

**Video 6.** Spatiotemporal dynamics in a two-dimensional simulation with Heaviside inhibition and homogeneous F-actin disassembly (Fig. 3D).

**Video 7.** Spatiotemporal dynamics in a two-dimensional simulation with Heaviside inhibition and heterogeneous F-actin disassembly (Fig. 3E).

**Video 8.** Spatiotemporal dynamics in a two-dimensional simulation with Cdk1-derived inhibition and homogeneous F-actin disassembly (Fig. 3F).

**Video 9.** Spatiotemporal dynamics on a spherical cortical shell with underlying regime (i) cytoplasmic dynamics (Fig. 4A).

**Video 10.** Spatiotemporal dynamics on a spherical cortical shell with underlying regime (ii) cytoplasmic dynamics (Fig. 4A).

**Video 11.** Spatiotemporal dynamics on a spherical cortical shell with underlying regime (iii) cytoplasmic dynamics (Fig. 4A).

**Video 12.** Spatiotemporal dynamics on a spherical cortical shell with underlying regime (iv) cytoplasmic dynamics (Fig. 4A).

**Video 13.** Effect of inhibition duration with homogeneous F-actin disassembly: short inhibition time (Supplementary Fig. S6A).

**Video 14.** Effect of inhibition duration with homogeneous F-actin disassembly: long inhibition time (Supplementary Fig. S6B).

**Video 15.** Effect of inhibition duration with heterogeneous F-actin disassembly: long inhibition time (Supplementary Fig. S6C).

**Video 16.** Spatiotemporal dynamics on a spherical cortical shell with underlying regime (i) dynamics and weak inhibition (large *s*) (Fig. 4C).

**Video 17.** Spatiotemporal dynamics on a spherical cortical shell with underlying regime (i) dynamics and strong inhibition (small *s*) (Fig. 4C).

**Video 18.** Experimental data from Bement et al. ***Bement et al. (2015)***; the bottom-left embryo is used for qualitative comparison with the model dynamics (Fig. 4E).

**Video 19.** Effect of nuclear position on cortical reactivation dynamics (Supplementary Fig. S7).

**Video 20.** Video with the merged normalised activity of Rho GTP (cyan) and F-actin (red) for *γ* = 1 (Fig. 4D).

## Data availability statement

The numerical code to reproduce the figures in this study is openly available on GitLab ***DiBS (2026)*** and archived with a persistent identifier in the KU Leuven Research Data Repository (RDR) ***Cebrián-Lacasa (2026)***. The generated data are openly available in the same repository ***Cebrián-Lacasa (2026)***.

## Notes

### Competing Interest Statement

The authors have declared no competing interest.

### Summary of Updates

This revised version includes minor clarifications to the Discussion, incorporating additional experimental evidence for nuclear Cyclin B-Cdk1 enrichment and compartment-specific activation dynamics. Figure captions and attribution statements have also been updated.

## References

Afanzar O, Buss GK, Stearns T, Ferrell J James E. The nucleus serves as the pacemaker for the cell cycle. eLife. 2020; 9:e59989. doi: 10.7554/eLife.59989.

Bement WM, Benink HA, Von Dassow G. A microtubule-dependent zone of active RhoA during cleavage plane specification. The Journal of cell biology. 2005; 170(1):91–101. doi: 10.1083/jcb.200501131.

Bement WM, Goryachev AB, Miller AL, von Dassow G. Patterning of the cell cortex by Rho GTPases. Nature Reviews Molecular Cell Biology. 2024; 25(4):290–308. doi: 10.1038/s41580-023-00682-z.

Bement WM, Leda M, Moe AM, Kita AM, Larson ME, Golding AE, Pfeuti C, Su KC, Miller AL, Goryachev AB, et al. Activator-inhibitor coupling between Rho signalling and actin assembly makes the cell cortex an excitable medium. Nature cell biology. 2015; 17(11):1471–1483. doi: 10.1038/ncb3251.

Beta C, Edelstein-Keshet L, Gov N, Yochelis A. From actin waves to mechanism and back: How theory aids biological understanding. eLife. 2023; 12:e87181. doi: 10.7554/eLife.87181.

Beta C, Kruse K. Intracellular Oscillations and Waves. Annual Review of Condensed Matter Physics. 2017; 8(1):239–264. doi: 10.1146/annurev-conmatphys-031016-025210.

Bischof J, Brand CA, Somogyi K, Májer I, Thome S, Mori M, Schwarz US, Lénárt P. A cdk1 gradient guides surface contraction waves in oocytes. Nature communications. 2017; 8(1):849. doi: 10.1038/s41467-017-00979-6.

Cebrián-Lacasa D, Leda M, Goryachev AB, Gelens L. Wave-driven phase wave patterns in a ring of FitzHugh-Nagumo oscillators. Physical Review E. 2024; 110(5):054208. doi: 10.1103/PhysRevE.110.054208.

Cebrián-Lacasa D, Replication Data for: "Geometry shapes cytoplasmic Cdk1 waves that drive cortical dynamics.". KU Leuven RDR; 2026. doi: 10.48804/RJ58AQ.

Cebrián-Lacasa D, Piñeros L, Vanderbeke A, Ruiz-Reynés D, Wouters T, Goryachev AB, Frolov N, Gelens L. Spiral waves speed up cell cycle oscillations in the frog cytoplasm. arXiv. 2024 12; doi: 10.48550/arXiv.2412.16094.

Chang JB, Ferrell Jr JE. Mitotic trigger waves and the spatial coordination of the Xenopus cell cycle. Nature. 2013; 500(7464):603–607. doi: 10.1038/nature12321.

Chomchai D, Leda M, Golding A, Von Dassow G, Bement WM, Goryachev AB. Rho GTPase dynamics distinguish between models of cortical excitability. Current Biology. 2025 Mar; 35(6):1414–1421.e4. doi: 10.1016/j.cub.2025.02.003.

Deneke VE, Di Talia S. Chemical waves in cell and developmental biology. Journal of Cell Biology. 2018 01; 217(4):1193–1204. doi: 10.1083/jcb.201701158.

Deneke V, Melbinger A, Vergassola M, Di Talia S. Waves of Cdk1 Activity in S Phase Synchronize the Cell Cycle in Drosophila Embryos. Developmental Cell. 2016; 38(4):399–412. doi: 10.1016/j.devcel.2016.07.023.

DiBS, 2026_Cebrián-Lacasa_Geometry. Gelens Lab, KU Leuven; 2026. https://gitlab.kuleuven.be/gelenslab/publications/coupledsystems_biomodels.

Foe VE, Alberts BM. Studies of nuclear and cytoplasmic behaviour during the five mitotic cycles that precede gastrulation in *Drosophila* embryogenesis. Journal of Cell Science. 1983; 61(1):31–70. doi: 10.1242/jcs.61.1.31.

Gavet O, Pines J. Activation of cyclin B1-Cdk1 synchronizes events in the nucleus and the cytoplasm at mitosis. The Journal of Cell Biology. 2010; 189(2):247–259. doi: 10.1083/jcb.200909144.

Gelens L, Anderson GA, Ferrell JE. Spatial trigger waves: positive feedback gets you a long way. Molecular Biology of the Cell. 2014; 25(22):3486–3493. doi: 10.1091/mbc.e14-08-1306.

Goryachev AB, Leda M. Autoactivation of small GTPases by the GEF–effector positive feedback modules. F1000Research. 2019 Sep; 8:1676. doi: 10.12688/f1000research.20003.1.

Goryachev AB, Leda M, Miller AL, Von Dassow G, Bement WM. How to make a static cytokinetic furrow out of traveling excitable waves. Small GTPases. 2016 Apr; 7(2):65–70. doi: 10.1080/21541248.2016.1168505.

Hara K. Cinematographic observation of "surface contraction waves" (SCW) during the early cleavage of axolotl eggs. Wilhelm Roux’ Archiv für Entwicklungsmechanik der Organismen. 1971; 167:183–186. doi: 10.1007/BF00577039.

Hara T, Abe M, Inoue H, Yu LR, Veenstra TD, Kang YH, Lee KS, Miki T. Cytokinesis regulator ECT2 changes its conformation through phosphorylation at Thr-341 in G2/M phase. Oncogene. 2006 1; 25:566–578. doi: 10.1038/sj.onc.1209078.

Kapadia N, Nurse P. Spatiotemporal orchestration of mitosis by cyclin-dependent kinase. Nature. 2025; 643:1391–1398. https://doi.org/10.1038/s41586-025-09172-y, doi: 10.1038/s41586-025-09172-y.

Klughammer N, Bischof J, Schnellbächer ND, Callegari A, Lénárt P, Schwarz US. Cytoplasmic flows in starfish oocytes are fully determined by cortical contractions. PLoS Computational Biology. 2018; 14(11):e1006588. doi: 10.1371/journal.pcbi.1006588.

Liu J, Burkart T, Ziepke A, Reinhard J, Chao YC, Tan TH, Swartz SZ, Frey E, Fakhri N. Light-induced cortical excitability reveals programmable shape dynamics in starfish oocytes. Nature Physics. 2025; 21:846–855. doi: 10.1038/s41567-025-02807-x.

Machacek M, Hodgson L, Welch C, Elliott H, Pertz O, Nalbant P, Abell A, Johnson GL, Hahn KM, Danuser G. Coordination of Rho GTPase activities during cell protrusion. Nature. 2009; 461(7260):99–103. doi: 10.1038/nature08242.

Maryu G, Yang Q. Nuclear-cytoplasmic compartmentalization of cyclin B1-Cdk1 promotes robust timing of mitotic events. Cell Reports. 2022; 41(13):111870. doi: 10.1016/j.celrep.2022.111870.

Michaud A, Leda M, Swider ZT, Kim S, He J, Landino J, Valley JR, Huisken J, Goryachev AB, von Dassow G, et al. A versatile cortical pattern-forming circuit based on Rho, F-actin, Ect2, and RGA-3/4. Journal of Cell Biology. 2022; 221(8):e202203017. doi: 10.1083/jcb.202203017.

Michaux JB, Robin FB, McFadden WM, Munro EM. Excitable RhoA dynamics drive pulsed contractions in the early *C. elegans* embryo. Journal of Cell Biology. 2018; 217(12):4230–4252. doi: 10.1083/jcb.201806161.

Miller PW, Stoop N, Dunkel J. Geometry of wave propagation on active deformable surfaces. Physical review letters. 2018; 120(26):268001. doi: 10.1103/PhysRevLett.120.268001.

Minc N, Piel M. Predicting division plane position and orientation. Trends in Cell Biology. 2012; 22(4):193–200. doi: 10.1016/j.tcb.2012.01.003.

Nalbant P, Hodgson L, Kraynov V, Toutchkine A, Hahn KM. Activation of endogenous Cdc42 visualized in living cells. Science. 2004; 305(5690):1615–1619. doi: 10.1126/science.1100367.

Niiya F, Tatsumoto T, Lee KS, Miki T. Phosphorylation of the cytokinesis regulator ECT2 at G2/M phase stimulates association of the mitotic kinase Plk1 and accumulation of GTP-bound RhoA. Oncogene. 2006 2; 25:827– 837. doi: 10.1038/sj.onc.1209124.

Nolet FE, Vandervelde A, Vanderbeke A, Piñeros L, Chang JB, Gelens L. Nuclei determine the spatial origin of mitotic waves. eLife. 2020; 9:e52868. doi: 10.7554/eLife.52868.

Novak B, Tyson JJ. Modeling the Cell Division Cycle: M-phase Trigger, Oscillations, and Size Control. Journal of Theoretical Biology. 1993 Nov; 165(1):101–134. doi: 10.1006/jtbi.1993.1179.

Piñeros L, Frolov N, Ruiz-Reynés D, Van Eynde A, Cavin-Meza G, Heald R, Gelens L. The nuclear-cytoplasmic ratio controls the cell cycle period in compartmentalized frog egg extract. Current Biology. 2025; 35(18):4426– 4441.e6. doi: 10.1016/j.cub.2025.07.024.

Puls O, Ruiz-Reynés D, Tavella F, Jin M, Kim Y, Gelens L, Yang Q. Spatial heterogeneity accelerates phase-to-trigger wave transitions in frog egg extracts. Nature Communications. 2024 12; 15:10455. doi: 10.1038/s41467-024-54752-7.

Rankin S, Kirschner MW. The surface contraction waves of Xenopus eggs reflect the metachronous cell-cycle state of the cytoplasm. Current Biology. 1997; 7(6):451–454. doi: 10.1016/S0960-9822(06)00192-8.

Rombouts J, Gelens L. Dynamic bistable switches enhance robustness and accuracy of cell cycle transitions. PLOS Computational Biology. 2021; 17(1):e1008231. doi: 10.1371/journal.pcbi.1008231.

Rossman KL, Der CJ, Sondek J. GEF means go: turning on RHO GTPases with guanine nucleotide-exchange factors. Nature reviews Molecular cell biology. 2005; 6(2):167–180. doi: 10.1038/nrm1587.

Santos SDM, Wollman R, Meyer T, Ferrell J James E. Spatial Positive Feedback at the Onset of Mitosis. Cell. 2012; 149(7):1500–1513. doi: 10.1016/j.cell.2012.05.028.

Satoh SK, Tsuchi A, Satoh R, Miyoshi H, Hamaguchi MS, Hamaguchi Y. The tension at the top of the animal pole decreases during meiotic cell division. PLoS ONE. 2013; 8(11):e79389. doi: 10.1371/journal.pone.0079389.

Shamipour S, Kardos R, Xue SL, Hof B, Hannezo E, Heisenberg CP. Bulk Actin Dynamics Drive Phase Segregation in Zebrafish Oocytes. Cell. 2019; 177(6):1463–1479.e18. doi: 10.1016/j.cell.2019.04.030.

Su KC, Takaki T, Petronczki M. Targeting of the RhoGEF Ect2 to the equatorial membrane controls cleavage furrow formation during cytokinesis. Developmental cell. 2011; 21(6):1104–1115. doi: 10.1016/j.devcel.2011.11.003.

Terasaki M, Okumura E, Hinkle B, Kishimoto T. Localization and dynamics of Cdc2-cyclin B during meiotic reinitiation in starfish oocytes. Molecular Biology of the Cell. 2003; 14(11):4685–4694. doi: 10.1091/mbc.e03-04-0249.

Tsai TYC, Theriot JA, Ferrell Jr JE. Changes in oscillatory dynamics in the cell cycle of early *Xenopus laevis* embryos. PLoS biology. 2014; 12(2):e1001788. doi: 10.1371/journal.pbio.1001788.

Wigbers MC, Tan TH, Brauns F, Liu J, Swartz SZ, Frey E, Fakhri N. A hierarchy of protein patterns robustly decodes cell shape information. Nature Physics. 2021; 17(5):578–584. doi: 10.1038/s41567-021-01164-9.

Yang Q, Ferrell Jr JE. The Cdk1-APC/C cell cycle oscillator circuit functions as a time-delayed, ultrasensitive switch. Nature cell biology. 2013; 15(5):519–525. doi: 10.1038/ncb2737.

Yin S, Li B, Feng XQ. Three-dimensional chiral morphodynamics of chemomechanical active shells. Proceedings of the National Academy of Sciences. 2022; 119(49):e2206159119. doi: 10.1073/pnas.2206159119.

